# Deciphering underground decarboxylase activity towards *N*_ε_-modified lysine derivatives in enterobacteria

**DOI:** 10.64898/2026.06.11.731642

**Authors:** Erica F. Aveta, Patroklos Vougioukas, Fei Qi, Judith Mehler, Kim Ina Behringer, Nicola Gericke, Marlene Walczak, Alexander P. Vallejo-Janeta, Thomas Blank, Michael Hellwig, Jürgen Lassak

## Abstract

Thermal food processing generates diverse compounds interacting with the gut microbiota. Despite their abundance, the microbial turnover of diet-borne *N*_ε_-modified lysine derivatives remains largely unexplored. We demonstrate that the enterobacterial ornithine decarboxylase SpeC degrades the prevalent advanced glycation end product *N*_ε_-carboxymethyllysine (CML) to carboxymethylcadaverine via an underground activity (∼4 molecules/enzyme/min). This promiscuity extends to additional *N*_ε_*-*modified lysine derivatives – namely fomylated (FmL), monomethylated (MML) and dimethylated (DML) lysine – yielding previously unknown biogenic amines (mono- and dimethylcadaverine, formylcadaverine). Functionally, SpeC enables *Escherichia coli* to utilize CML as a sole nitrogen source. In specific strains, this metabolism reinforces pH-stress responses, supporting survival under mild acidic conditions typical for the colon. Furthermore, SpeC orthologs are widespread across human gut genomes, correlating with geography, diet, and disease. Together, these findings suggest a potential diet-microbiome communication axis, linking the intake of modified dietary chemicals to microbial physiology and hypothesized host impacts.

## 1. Introduction

Non-canonical amino acids are frequently found in our regular diet (Hellwig et al., 2024). Thermal food processing further promotes glycation and hence the enrichment of so-called AGEs which are formed during the late stage of the Maillard reaction (Singh et al., 2001). Among the diverse array of glycation products, CML serves as a prominent and widely recognized marker for dietary AGEs (Hellwig et al., 2024). CML is generated through parallel chemical mechanisms, including both oxidation of Schiff bases and Amadori products as well as the reaction with glyoxal that may originate from Maillard reaction or lipid peroxidation (Delgado-Andrade, 2016). Up to 24.6 mg of CML are ingested daily (Delgado-Andrade, 2016; Mark et al., 2014). CML is incompletely released from dietary proteins during digestion and in parts transported across the intestinal epithelium (Delgado-Andrade, 2016; Hellwig et al., 2011). Notably, this permeability increases in an age- and microbiota-dependent manner, leading to increased CML levels in serum and brain (Mossad et al., 2022; Thevaranjan et al., 2017). This in turn is correlated with the impairment of the function of microglia, the primary immune cells of the central nervous system. Here, CML causes a burst of reactive oxygen species and impairs mitochondrial activity and adenosine 5’-triphosphate (ATP) storage (Mossad et al., 2022). Most interestingly, the gut microbiota can not only promote an increase of CML levels in humans, but is also capable of degrading it (Delgado-Andrade, 2016; Hellwig et al., 2015; Hellwig et al., 2011). The biogenic amine of CML – CM-Cad – is the major reaction product in CML degrading gut bacteria including *E. coli* (Bui et al., 2019; Hellwig et al., 2019). However, no enzyme catalyzing the decarboxylation has been reported to date. In this study, we identify the ornithine decarboxylase SpeC as the enzyme responsible for the decarboxylation of CML and other *N*_ε_-modified lysine derivatives in Enterobacteria. Due to the low substrate turnover rates we postulate that this degradative capacity represents a classic example of underground metabolism (D’Ari & Casadesús, 1998; Rosenberg & Commichau, 2019). In molecular and evolutionary systems biology, this paradigm describes the collection of latent, promiscuous enzyme side activities within a cell. While enzymes face evolutionary pressure to optimize their primary physiological functions, their inherent structural flexibility allows them to process structurally related, non-canonical substrates at very low catalytic rates. These inefficient, underground activities serve as a vital evolutionary reservoir, enabling microorganisms to rapidly adapt to environmental stressors or novel nutritional inputs, such as the influx of dietary AGEs. We demonstrate that this underground activity enables *E. coli* to utilize CML as a nitrogen source and, in certain strains, confers a competitive advantage by reinforcing the acid stress response. Furthermore, bioinformatic analyses reveal that SpeC orthologs are widespread in the human gut microbiome and correlate with westernized diets. Finally, we show that the resulting metabolite, CM-Cad, is readily internalized by microglia, suggesting a potential link between dietary metabolites and host innate immunity. To provide a clear overview of our multidisciplinary approach, we have included a conceptual study design (Scheme 1), which maps our core hypotheses – ranging from the initial enzyme discovery to the assessment of physiological and ecological relevance – thereby including the corresponding experimental strategies and measured indicators.

**Scheme 1:**
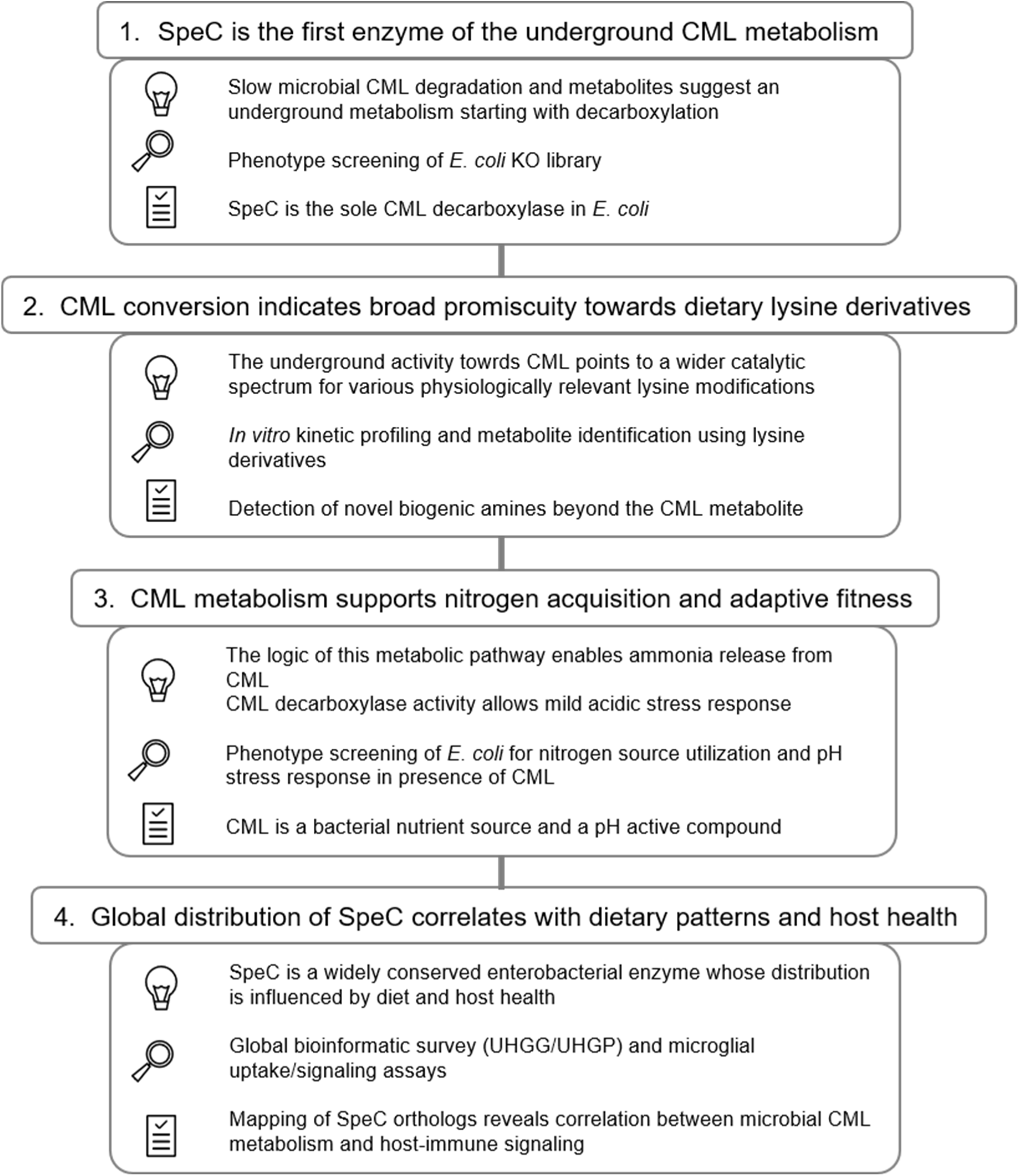
Study design

## 2. Materials and Methods

### 2.1. Media and growth conditions

*E. coli* was cultivated in Lysogeny Broth (LB), Super Optimal Broth (SOB), or M9 minimal medium (33.7 mM Na_2_HPO_4_, 22.0 mM K_2_HPO_4_, 8.55 mM NaCl, 1 mM MgSO_4_, 0.3 mM CaCl_2_, 1 µg/L biotin and thiamine, 13.4 mM ethylenediaminetetraacetic acid (EDTA), 3.1 mM FeCl_3_, 0.62 mM ZnCl₂, 76 µM CuCl₂, 42 µM CoCl_2_, 162 µM H_3_BO_3_, 8.1 µM MnCl_2_) (Bertani, 1951; Hanahan, 1983; Miller, 1972). Unless otherwise stated, M9 medium was supplemented with 10 mM NH_4_Cl as the nitrogen source and 20 mM glucose or glycerol as the carbon source. Where necessary antibiotics were added at the following concentrations: 50 μg/mL kanamycin sulfate, 34 μg/mL chloramphenicol. Cultures were incubated at 37°C. To induce expression from pET or pBAD vectors, 1 mM isopropyl β-D-thiogalactoside (IPTG) or 0.2% (w/v) L-(+)-arabinose was added, respectively, when cells entered logarithmic growth phase.

For long-term storage, strains were cryopreserved at -80 °C in glycerol freezing medium (87% glycerol, 1 M KCl, 1 M NaCl, 0.2 M MgSO₄) at a 1:3 ratio. Prior to physiological assays in M9 medium, cells from rich medium precultures were washed twice (15,000 × g, 1.5 min) with M9 medium lacking carbon and nitrogen to prevent nutrient carryover. Growth was monitored by measuring optical density at 600 nm (OD_600_) using an Ultrospec 2100pro spectrophotometer (Amersham Biosciences) or a thermostatic plate reader (Tecan Infinite 500 or Spark 20M).

### 2.2. Genetic manipulation

#### 2.2.1. Cloning and plasmid construction

Chemically competent *E. coli* cells were prepared and transformed using the CaCl₂ method according to (Dagert & Ehrlich, 1979). Coding sequences (CDS) for *speC* and *speF* were amplified from genomic DNA of *E. coli* MG1655 – extracted according to (Pospiech & Neumann, 1995) – using Q5® HiFi DNA polymerase (New England Biolabs). Orthologs from *Vibrio cholerae* O1 biovar El Tor N16961, *Salmonella enterica* subsp*. enterica* serovar Typhimurium 14028s, and *Yersinia enterocolitica* NCTC12982 were similarly amplified. Corresponding strains, plasmids and primers are listed in the tables S4-6.

For purification and western blot detection, genes were cloned into pET-SUMO (Thermo Fisher) or pBAD33 (fused with a C-terminal 6xHis-tag coding sequence). For pBAD33 constructs, a synthetic ribosome binding site was designed using the RBS calculator (Cetnar & Salis, 2020; Reis & Salis, 2020). Restriction digestion and ligation (T4 ligase) were performed according to New England Biolabs protocols. PCR products and vectors were purified using the HiYield® PCR Clean-Up & Gel-Extraction kit (Süd-Laborbedarf). Constructs were maintained in *E. coli* DH5αλpir (Taylor et al., 1993). Plasmid DNA was isolated using the HiYield® Plasmid Mini Kit (Süd-Laborbedarf) and quantified spectrophotometrically.

#### 2.2.2 **Deletion of** *speC* **in** *E. coli* **Nissle 1917**

The *speC* gene was deleted in *E. coli* Nissle 1917 via homologous recombination using the Quick & Easy *E. coli* Gene Deletion Kit (Gene Bridges GmbH) (Baba et al., 2006). The CDS of *speC* was replaced with a kanamycin resistance cassette flanked by FLP recognition sites. Strains, Primers and plasmids (pRed-ET, pKD13) are listed in tables S4-S6.

#### 2.2.3 Sequencing

All plasmid constructs and mutant strains were verified by Sanger sequencing at the Genomics Service Unit (LMU Munich).

### 2.3. *In vivo* degradation and physiological assaysh

#### 2.3.1. AGE degradation and complementation assay

To assess CML degradation capabilities, *E. coli* BW25113 wild type and single-gene deletion mutants (Δ*cadA*, Δ*ldcC*, Δ*gadA*, Δ*gadB*, Δ*adiA*, Δ*speA*, Δ*speC*, Δ*speF*, Δ*lysA*) were precultured in M9 glucose medium (supplemented with 50 µM kanamycin sulfate or L-lysine where appropriate). Washed cells were inoculated to an OD_600_ of 0.3 in M9 medium containing 250 µM CML (but no NH₄Cl) and incubated at 37°C and grown under microaerophilic conditions. Samples were collected at 0, 4, 24, 48, and 72 h and stored at -20 °C for HPLC-MS analysis.

For complementation, *E. coli* Δ*speC* harboring pBAD33-speC or an empty vector were grown in M9 glycerol medium with antibiotics. Expression was induced with 0.2% L-(+)-arabinose at OD_600_ 0.3–0.4. After 4 h of induction, 250 µM CML was added. Samples were taken and after 24 h incubation.

Protein expression was verified by SDS-PAGE and Western Blot. Samples normalized to an OD_600_ of 10 were separated and blotted onto nitrocellulose membranes (0.45 µm). Proteins were detected using anti-His tag antibodies (Thermo Fisher or Abcam) and alkaline phosphatase-conjugated secondary antibodies (Rockland), visualized with NBT/BCIP solution.

#### 2.3.2. Utilization of AGEs as a nitrogen source

*E. coli* BW25113 and JW5482 (Δ*speC*) were grown overnight, washed, and inoculated (OD_600_ 0.01) into 96-well plates containing M9 glucose medium with distinct nitrogen sources: 10 mM CML, L-ornithine, L-lysine, NH_4_Cl, or no nitrogen. Growth was recorded every 30 min for 48 h at 37°C. Maximum OD increase was calculated from three biological replicates.

#### 2.3.3. Acid stress response assay

Acid stress response was monitored using a colorimetric assay based on bromocresol purple (BCP) (Møller, 1955). Strains (including *E. coli* K-12, Nissle 1917, and clinical isolates) were washed and inoculated (OD_600_ 0.01) into decarboxylase medium (0.5% tryptone, 0.3% yeast extract, 0.1% glucose, 37 µM BCP, 20 µM pyridoxal phosphate, pH 5.0) supplemented with 10 mM L-ornithine, L-lysine, CML, or no amino acid as control condition. To determine the ratiometric pH value, absorbance was measured at 432 nm (Abs_432_) and 600 nm (Abs_600_), corresponding to the absorption maxima of the protonated (acidic) and deprotonated (basic) forms of bromocresol purple, respectively. The ratio of Abs_600_/Abs_432_ was calculated after correcting for cell turbidity (measured in BCP-free controls) and background absorbance.

### 2.4. *In vitro* **AGE degradation**

#### 2.4.1 Protein expression and purification

*E. coli* BL21(DE3) transformed with pET-SUMO constructs encoding SpeC or SpeF variants were grown in LB/kanamycin to OD_600_ 0.5 at 37 °C. Expression was induced by adding 1 mM IPTG followed by an overnight incubation at 18 °C. Cells were harvested, resuspended in lysis buffer (50 mM Tris, 300 mM NaCl, 1,4-dithiothreitol (DTT), 1 mM phenylmethylsulfonyl fluoride (PMSF), pH 7.6), and disrupted using a high-pressure homogenizer (1.35 kbar) followed by sonication.

Proteins were purified from the cytosolic fraction via Ni^2+^-affinity chromatography (Ni-NTA Agarose, Qiagen). After washing (20 mM imidazole), proteins were eluted with 250 mM imidazole. The SUMO-tag was cleaved by Ulp1 protease digestion during dialysis against lysis buffer and subsequently removed by reverse Ni^2+^-affinity chromatography. Protein purity and concentration were assessed by SDS-PAGE and OD_280_ measurements using a NanoDrop spectrophotometer (Thermo Fisher Scientific), respectively.

#### 2.4.2. Enzyme kinetics and substrate specificity

Enzymatic assays were performed in reaction buffer (50 mM Tris, 300 mM NaCl, 0.1 mM GTP, 0.1 mM pyridoxal phosphate, pH 7.0) at 37 °C.

For substrate specificity, 50 or 500 nM enzyme was incubated with 10 mM substrate (L-ornithine, L-lysine, L-arginine, CML, CEL, AcL, MML, DML, TML, FmL). Reactions were stopped with 200 mM Na₂CO₃ at various time points (up to 24 h). Substrate and product concentrations were quantified by HPLC.

Kinetic parameters for *E. coli* SpeC and SpeF were determined using substrate concentrations ranging from 0.05 to 10 mM. Initial velocities were calculated from linear product formation phases (0–5 min for ornithine; 0–48 h for AGEs). Data were fitted using SigmaPlot 12 to obtain K_M_ and v*_max_*; the turnover rate (k*_cat_*) was subsequently calculated based on the molecular weight of the enzyme.

#### 2.4.3. Sequence alignment and active site comparison

Multiple sequence alignment of the ornithine decarboxylases used for this study was performed with Clustal Omega (Madeira et al., 2024). The amino acid sequences were retrieved for SpeC and SpeF of each strain from Uniprot or NCBI and their accession numbers are the following: *E. coli* SpeC (P21169) and SpeF (P24169), *S. enterica* Typhimurium SpeC (A0A0F6B6L9) and SpeF (A0A0F6AYK0), *Y. enterocolitica* SpeC (NCBI, VTP82757.1) and SpeF (NCBI, VTP87433.1), *V. cholerae* SpeF (NCBI, AAF96957). The alignment included the reference ornithine decarboxylase OdcI from *Lactobacillus* 30A (Uniprot ID P43099) (Momany et al., 1995) which is not part of the *in vitro* study. The comparison of the structures and models, retrieved from PDB or SWISS-MODEL in the respective Uniprot page, was done using ChimeraX (Meng et al., 2023).

### 2.5. Analytical methods

#### 2.5.1. Synthesis of carboxymethylated compounds as standards

CML, CEL and CM-Cad and were synthesized as previously described (Hellwig et al., 2019; Hellwig et al., 2011). ^1^H NMR spectra were obtained on a Bruker Avance II device (300 MHz), and ^13^C spectra on a Bruker Avance 600 device (150 MHz). Mass-spectrometric data were obtained by tandem mass spectrometric analysis with the analytical system described in chapter 2.5.2, while high-resolution data were obtained on a Bruker Impact II device. Analytical data are as follows. ^1^H and ^13^C NMR spectra with the atom numbering are included in Figures S1-S4, as well as the mass spectra of CML and CM-Cad (Figure S5).

(CM-Cad). ^1^H NMR (300 MHz, D_2_O) δ [ppm]: 1.39 (m, 2H, H-3); 1.66 (m, 4H, H-2, H-4); 2.94 (t, 2H, J = 6.9 Hz, H-1); 3.05 (t, 2H, J = 7.6 Hz, H-5); 3.83 (s, 2H, H-1’). ^13^C NMR (150 MHz, D_2_O) δ [ppm]: 22.8 (2 CH_2_, C-3), 25.0 (CH_2_, C-4), 26.3 (CH_2_, C-2), 39.2 (CH_2_, C-1), 47.3 (2 CH_2_, C-1’, C-5), 168.8 (C_i_, C-2’). HR-MS: calcd., 161.129; found, 161.123. HPLC-MS/MS: t_R_, 9.0 min; fragmentation (80 V, 20 eV) of [M + H]^+^ (*m/z* 161) 86 (100), 98 (61), 69 (18). Anal. Calcd for C_7_H_16_N_2_O_2_ (MW = 160.21): C, 52.48; H, 10.07; N, 17.48; C/N, 3.00. Found: C, 19.93; H, 4.68; N, 6.63; C/N, 3.01. Content 37.9 % based on nitrogen. Yield: 550.2 mg (33 %).

(CML). ^1^H NMR (300 MHz, D_2_O), δ [ppm]: 1.45 (m, 2H, H-4); 1.70 (m, 2H, H-5); 1.90 (m, 2H, H-3); 3.04 (t, 2H, H-6, J = 7.8); 3.84 (s, 2H, H-1’) 3.98 (t, 1H, J = 6.3 Hz, H-2). ^13^C NMR (150 MHz, D_2_O) δ [ppm]: 21.5 (CH_2_, C-4), 25.0 (CH_2_, C-5), 29.3 (CH_2_, C-3), 47.1 (CH_2_, C-6), 47.4 (CH_2_, C-1’), 52.7 (CH, C-2), 169.2 (C_i_, C-2’), 172.1 (C_i_, C-1). HR-MS: calcd., 205.119; found, 205.111. HPLC-MS/MS: t_R_, 7.4 min; fragmentation (80 V, 20 eV) of [M + H]^+^ (*m/z* 205) 84 (100), 130 (6), 142 (5). Anal. Calcd for C_8_H_16_N_2_O_4_ (MW = 204.22): C, 47.05; H, 7.90; N, 13.72; C/N, 3.43. Found: C, 30.95; H, 6.28; N, 9.10; C/N, 3.40. Content 66.3 % based on nitrogen. Yield: 250.4 mg (20 %).

#### 2.5.2. Quantification of metabolites via HPLC-UV and HPLC-MS

Amino acids and biogenic amines were qualitatively analyzed and/or quantified after dansylation. Samples (10-30 µL) were diluted to a volume of 100 µL with water, and 150 µL of 0.1 M sodium carbonate solution as well as 200 µL of a solution of dansyl chloride in acetone (0.5%, w/v) were added. After incubation in a water bath (40 °C, 1 h), 10 µL of 3 M HCl was added. After shaking, 540 µL of 0.1% aqueous formic acid was added. The solutions were centrifuged (10,000 × g, 10 min, 4 °C), and the supernatants were transferred to HPLC vials. An aliquot (20 µL) was injected into a UHPLC system from VWR (Darmstadt, Germany) equipped with a diode array detector (DAD 6430) that operated at λ = 254 nm, while spectra were recorded between 200 and 400 nm. As solvent A, 0.1% formic acid in water was used, and 0.1% formic acid in a mixture of acetonitrile and water (90/10, v/v) as solvent B. Separations were performed at 40 °C on a C-18 column (Nucleodur-100 endcapped, 250 × 4.6 mm, 5 µm, Macherey-Nagel). The flow rate was 1 mL/min, and a gradient was used for all dansylated amino acids (0 min, 20% B; 17 min, 67% B; 25 min, 100% B; 29 min, 100% B; 30 min, 20% B; 34 min, 20% B), except dansylated DML and TML (0 min, 10% B; 5 min, 10% B; 15 min, 20% B; 20 min, 40% B; 25 min, 40% B; 26 min, 10% B; 31 min, 10% B). For dansylated FmL and AcL, a further gradient was applied (0 min, 20% B; 29 min, 67% B; 30 min, 20% B; 34 min, 20% B).

For ESI-TOF analysis, 20 µL of the dansylated samples were diluted with 180 µL solvent A, membrane filtered (0.2 µm) and transferred to HPLC vials. An aliquot (1 µL) was injected into a UHPLC system Infinity 1290 (Agilent) consisting of a binary pump with an integrated degasser, an autosampler (G4226A), and a UV-detector (G4212A) that was connected to the mass spectrometer TIMS-TOF (Bruker). UV detection was performed at λ = 254 nm, and spectra were recorded between 210 and 450 nm (increment, 1 nm). The same solvents as above were used. Separations were performed at 40°C on a stainless-steel column filled with the material Eurospher-II 100 (50 mm × 2.0 mm, 3 µm, Knauer) The flow rate was 0.2 mL/min, and a gradient was used (0 min, 20% B; 17 min, 67% B; 25 min, 100% B; 30 min, 100% B; 31 min, 20% B; 37 min, 20% B). The mass spectrometer operated in the positive scan mode (*m/z* 20-1300; scan time, 500 ms; dry gas flow, 10 L nitrogen/min; dry temperature, 220 °C; nebulizer pressure, 2.2 bar; capillary voltage, 4500 V).

For quantification by tandem mass spectrometric analysis, 5 µL of sample was mixed with either 5 µL 5 mM HCl or 5 µL of a standard solution (1 mM CML, 0.1 mM CM-CAD, 1 mM ornithine, and 0.1 mM putrescine) as well as 90 µL of water. After centrifugation (10,000 × g, 10 min, 4 °C), the sample was analyzed on an Agilent 1200 Series system connected to the Triple Quad 6410 tandem mass spectrometer. The injection volume was 5 µL. Separation was performed on a Zorbax 100 SB-C18 column (2 x 50 mm, Agilent) at 35°C. At a flow rate of 0.25 mL/min, a gradient was formed from the solvents A (10 mM nonafluoropentanoic acid (NFPA) in water) and B (10 mM NFPA in acetonitrile) (0 min, 10% B; 15 min, 66% B; 19 min, 66% B; 20 min, 10% B; 25 min, 10% B).

The following settings were applied at the mass spectrometer: capillary voltage, 4000 V; source temperature, 350°C; gas flow, 11 L nitrogen/min; nebulizer pressure, 35 psi. The spectrometer operated in the scan mode (*m/z* 20-1,300; scan time, 500 ms), or in the MRM mode with the parameters compiled in table S3. Transitions were recorded between 2 and 13 min.

### 2.6. Bioinformatic analysis of the distribution of SpeC degrading bacteria in the human gut microbiome

Orthologs of *E. coli* SpeC: using the OMA database (Altenhoff et al., 2021), we obtained 739 orthologous proteins of *E. coli* SpeC from 611 strains representing 387 species. Taxonomic information for these species was obtained from the NCBI taxonomy database (Schoch et al., 2020), and was visualized using the *ggtree* package in the R environment (Dimaano et al., 2024). Each orthologous protein sequence was then aligned to the amino acid sequence of *E. coli* SpeC using the *needleall* program from the EMBOSS package (Rice et al., 2000). The resulting pairwise sequence alignments were used to calculate the sequence identities between the orthologous proteins and *E. coli* SpeC.

Occurrence of CML-degrading bacteria in the human gut microbiota: We investigated this using the Unified Human Gastrointestinal Genome (UHGG) and Protein (UHGP) collections, which contain 630,333,479 proteins from 289,232 prokaryotic genomes clustered into 4,744 species (Almeida et al., 2021). We downloaded the 95,303,635 protein clusters at 100% sequence identity (UHGP-100) from the collections. The representative sequences of each cluster were aligned to the *E. coli* SpeC amino acid sequence using the *needleall* program from the EMBOSS package and the resulted pairwise sequence alignments were used to calculate the sequence identity (Rice et al., 2000). For each protein cluster, if its representative sequence showed ≥ 70.4% sequence identity with *E. coli* SpeC (defined based on the sequence identities between SpeC orthologs of Enterobacterales and *E. coli* SpeC from the above orthology analysis), all members of this cluster were identified as capable of degrading CML, and their corresponding genomes were identified as CML-degrading. Using this procedure, we identified 10,330 proteins from 1,476 clusters and 10,252 genomes from 89 species capable of degrading CML.

Metadata of the human gut microbiota samples: The geographic information of the human gut microbiota samples was from the UHGG collection. The metadata for the lifestyle, age category and disease of the samples were derived from the *curatedMetagenomicData* package (version 3.8.0, in the R environment) (Pasolli et al., 2017). The microbiota samples between the UHGG and the *curatedMetagenomicData* package were mapped using their accession numbers. The metadata for the microbiota samples of hunter-gatherers were manually curated from (Obregon-Tito et al., 2015; Rampelli et al., 2015).

### 2.7. Microglial cells treatment

#### 2.7.1. Isolation of glial cells

All animal experiments were approved by the local administration (Regierungspräsidium Freiburg, approval number X-2402a issued Febr. 1, 2024) and were performed in accordance with the respective national, federal and institutional regulations. Primary cultures of microglia and astrocytes were prepared from neonatal (P0–P3) C57BL/6N mice (CEMT, University Medical Center Freiburg) as previously described (Blank & Prinz, 2013). The mice were euthanized by decapitation in accordance with animal care guidelines. Brains were removed under aseptic conditions and placed in ice-cold Hanks’ Balanced Salt Solution (HBSS, 14170088, Gibco). Under a stereomicroscope, meninges were carefully removed, and cortices were dissected and transferred to a 6-well plate containing ice-cold mixed glial medium formulated with DMEM/F12 (11320033, Gibco) supplemented with 20% heat-inactivated fetal bovine serum (hi-FBS, A5256701, Gibco) and 1% penicillin/streptomycin (P/S, P0781, Sigma-Aldrich).

Cortical tissue was mechanically dissociated by gently pipetting. The cell suspension was transferred to T-25 flasks containing 1 mL of mixed glial medium and maintained at 37 °C in a humidified atmosphere with 5% CO_2_. Cells were grown for 10 days, followed by 2-4 days of culture in mixed glial medium supplemented with 20 ng/mL recombinant murine colony-stimulating factor (M-CSF, 315-02-10UG, PeproTech) until confluency was reached.

#### 2.7.2. Microglial cells harvest and seeding experiments

Upon reaching confluency, microglial cells were harvested from the mixed glial culture by mechanical detachment placing the T-25 flasks on a shaking incubator at 37 °C and 180 rpm for 1 h. The supernatant containing detached microglia was collected and centrifuged at 400 × g for 5 min at room temperature. The cell pellet was resuspended in microglial medium (DMEM/F12 supplemented with 10% hi-FBS, 1% P/S, and 20 ng/mL M-CSF).

The microglial cell suspension was seeded onto poly-D-lysine (PDL, A3890401, Gibco)-coated T-25 flasks containing 5 mL of fresh microglial medium. Flasks were gently swirled and incubated for 30 min at room temperature under sterile conditions to ensure even cell distribution, followed by 1 h incubation at 37 °C. The medium was then completely replaced with fresh microglial medium to remove non-adhered cells. Cells were cultured until confluency with medium replacement every 2-3 days.

Confluent microglial cultures were detached by trypsinization with 0.25% trypsin-EDTA, incubated at 37 °C for 5 min. Trypsinization was quenched by adding 4 mL of microglial medium, and cells were completely detached by gentle pipetting. The cells were pelleted by centrifugation and the resulting cell pellet was resuspended. Microglial cells were seeded into 8-well chamber slides (154534, Thermo Scientific) at a density of 5×10^4^ cells per well in 300 µL of microglial medium. Cells were allowed to adhere for 24 h at 37 °C before treatment with CML and CM-Cad.

#### 2.7.3. Uptake of CML and CM-Cad in murine microglial cells

Microglial cells were treated with 100 µM CML or 100 µM CM-Cad for 7 days. The medium was changed every 2 days containing the same CML or CM-Cad concentrations. Control cells received medium changes at the same intervals without treatment compounds. One day after the final treatment, cells were fixed with 4% paraformaldehyde (PFA) in phosphate-buffered saline (PBS) for 15 min at room temperature, followed by permeabilization with PBS containing 0.1% Triton X-100 for 5 min. Non-specific binding was blocked using blocking buffer composed of 1X PBS, 5% bovine serum albumin (BSA), 0.5% Triton X-100, and 0.02% sodium azide. Primary antibodies were diluted in blocking buffer as follows: rabbit anti-Iba1 (1:750, Abcam, ab178846) and mouse anti-CML (1:200, Abcam, ab125145). Cells were incubated with primary antibodies overnight at 4 °C, then with secondary antibodies for 1 h at room temperature: Alexa Fluor 488-conjugated donkey anti-rabbit IgG (1:500, Invitrogen, A-21206) and Alexa Fluor 647-conjugated donkey anti-mouse IgG (1:500, Invitrogen, A-31571). Nuclei were counterstained with DAPI (1:10,000) for 10 min. Slides were mounted with Mowiol and stored at 4 °C until imaging. Fluorescence images were acquired using a Leica DMi8 microscope at 20× magnification. For each treatment condition, multiple regions of interest were captured to ensure representative sampling. Quantification was performed using ImageJ, and values were reported as CML-positive cells relative to the total number of cells. Data were presented as mean ± s.e.m. Statistical analyses were conducted using GraphPad Prism 10. One-way Brown-Forsythe and Welch ANOVA was performed, followed by Dunnett’s T3 multiple comparison test for post hoc pairwise comparisons. Statistical significance was set at α = 0.05, with significance levels denoted as: \**P* < 0.05, \*\**P* < 0.01, \*\*\**P* < 0.001, and \*\*\*\**P* < 0.0001.

### 2.8 Statistical analysis

Experimental data are generally expressed as the mean ± standard deviation (SD) of independent biological replicates, unless otherwise stated (e.g., murine microglial assays are presented as the mean ± standard error of the mean (SEM)). Statistical evaluations were performed using appropriate software, including GraphPad Prism and the R environment. Depending on the experimental design, data were analyzed using Student’s t-test, analysis of variance (ANOVA), or Fisher’s Exact test. Statistical significance was defined as p < 0.05 (*p < 0.05, **p < 0.01, ***p < 0.001, ****p < 0.0001). Detailed information regarding the specific statistical tests applied, the exact number of replicates (n), and the respective measures of dispersion for each experiment are detailed in the figure legends.

## 3. Results and discussion

To identify enzymes responsible for the conversion of CML to the main degradation product CM-Cad (Fig. 1a) (Hellwig et al., 2019), we screened for *E. coli* proteins with known decarboxylase activity using the EcoCyc database (Table S1) (Keseler et al., 2021; Sayers et al., 2022). We ranked the enzymes according to the chemical similarity of their cognate substrates to CML, chose nine candidates and tested corresponding *E. coli* deletion strains from the Keio collection (Baba et al., 2006) on their ability to degrade CML to CM-Cad (Fig. 1b). All strains were able to decarboxylate CML except the mutant lacking the ornithine decarboxylase SpeC (Δ*speC*). We took advantage of the CM-Cad null mutant Δ*speC*, which was transformed with arabinose inducible copies of SpeC. CM-Cad production in Δ*speC* could readily be restored by ectopic expression of *speC*, in fact, 40-60% of CML was converted to CM-Cad, compared to the ∼5% in the wild type (Figs. 1b, c). This result confirms SpeC as an important player in CML degradation.

**Fig. 1.**
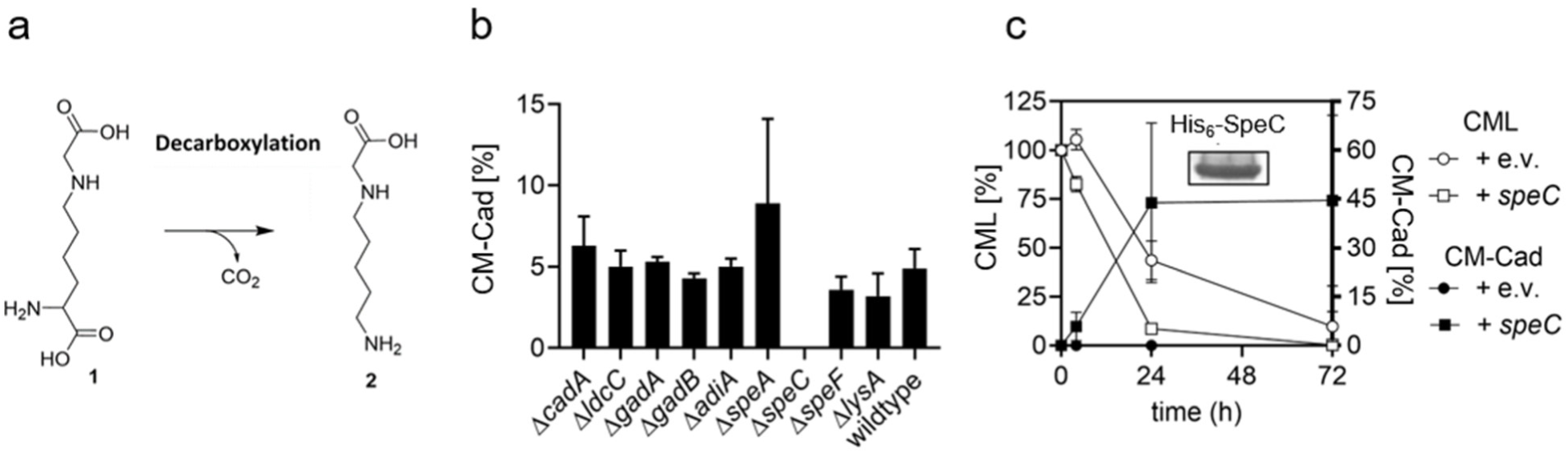
Discovery of a CML decarboxylase in *E. coli*. a) Degradation of CML **1** to CM-Cad **2**, b) CM-Cad formation in isogenic *E. coli* K-12 BW25113 decarboxylase mutant strains. Cells were incubated with 250 µM of CML for 24 h in M9 minimal medium. CM-Cad levels were determined after dansyl-precolumn derivatization by HPLC-UV and are given in % relative to initial CML. The results are represented as mean and standard deviation of two independent biological replicates c) Complementation analysis of the *E. coli* Δ*speC* deletion strain. Δ*speC* cells either containing an empty vector (e.v.) or ectopically overexpressing the ornithine decarboxylase *speC*, from an arabinose inducible promoter were incubated with 250 µM of CML for 72 h in M9 minimal medium. CML and CM-Cad levels were determined after dansyl-precolumn derivatization by HPLC-UV and are given in % relative to initial CML. The results are represented as mean and standard deviation of three independent biological replicates.

As our *in vivo* approach gave only semiquantitative data, we next assessed the enzymatic properties of SpeC *in vitro*. Notably, SpeC belongs to the constitutively expressed decarboxylases, whereas others are inducible and thus might have been overlooked in our initial screening (Fig. 1c) (Kanjee, Gutsche, Ramachandran, et al., 2011). The latter category includes, most interestingly, the SpeC paralog SpeF (with a sequence identity of 66.8%) (Kanjee, Gutsche, Ramachandran, et al., 2011). Accordingly, SpeF was tested alongside SpeC. An initial time course experiment was set up for both enzymes in presence of 10 mM CML (Fig. 2a) and ornithine (Figs. S6 and S7). As *in vivo*, SpeC degraded CML to CM-Cad (Fig. 2a), while SpeF did not show any detectable activity. It is important to note here that while ornithine decarboxylases are found in both Fold Type I and Fold Type III of pyridoxal 5’-phosphate (PLP)-dependent enzymes, both SpeC and SpeF belong to the prokaryotic ornithine decarboxylases of Fold Type I (Deng et al., 2010; Kanjee, Gutsche, Alexopoulos, et al., 2011; Phillips et al., 2019). Unlike Fold Type III ornithine decarboxylases, which utilize a β/α-barrel architecture with a defined specificity helix that determines substrate specificity, Fold Type I decarboxylases lack such clearly defined structural elements. Although specific active site residues have been hypothesized to act as substrate gating mechanisms, these are strictly conserved between SpeC and SpeF (Fig. S8). Sequence alignments reveal only two notable differences: An N-terminal gap (between amino acids 80 and 100) in the SpeC orthologs of *E. coli* and *S. enterica* as well as a C-terminal extension in their respective SpeF counterparts (Figs. S8 and S9). However, since both structural variations are absent in the functionally active SpeC ortholog of *Y. enterocolitica* (Fig. 4a), they likely do not account for the observed substrate promiscuity.

**Fig. 2.**
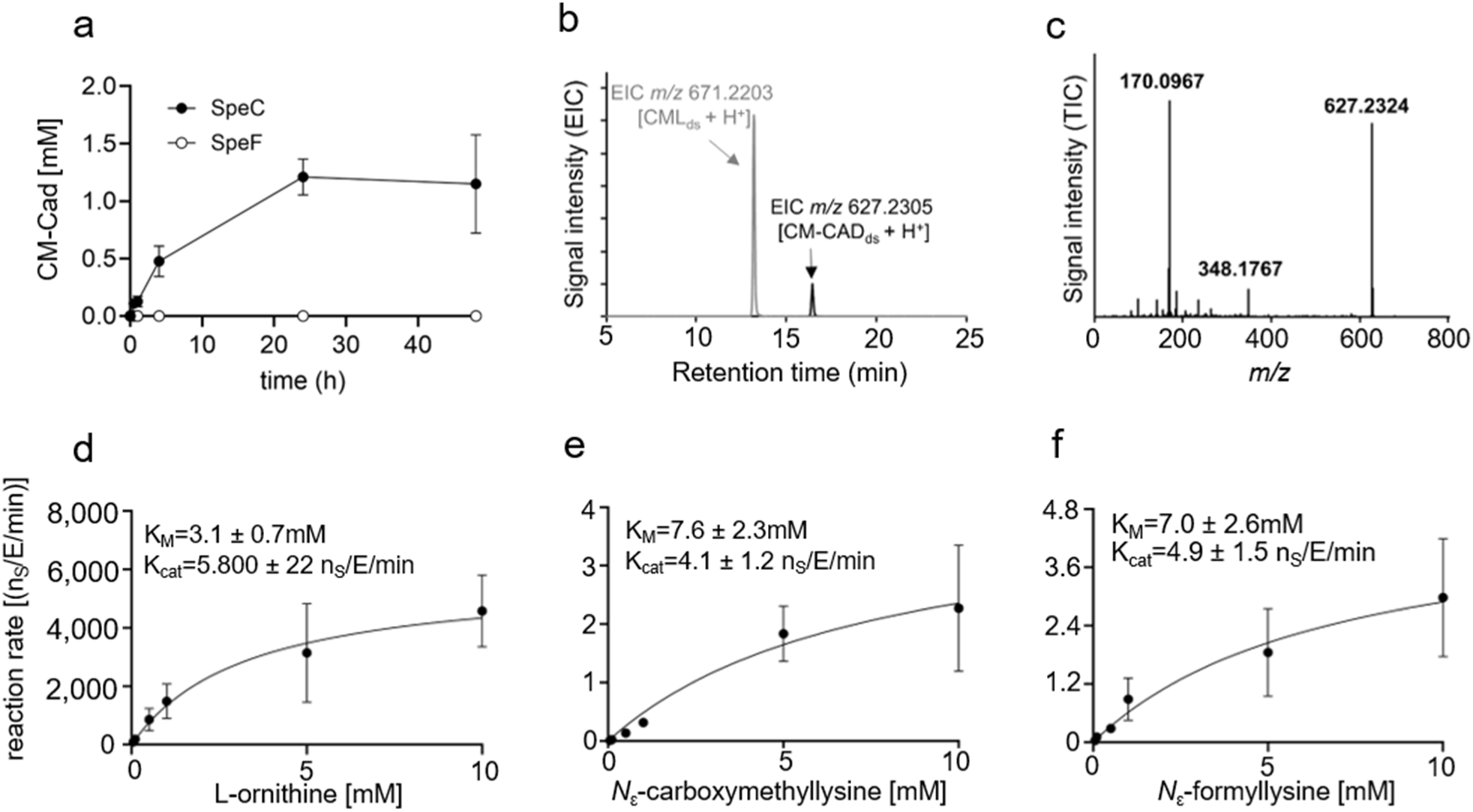
*In vitro* characterization of SpeC. a) *In vitro* CM-Cad formation over time resulting from the degradation of CML, in presence of SpeC or SpeF purified from *E. coli* K-12. CM-Cad levels were determined by HPLC-UV or HPLC-MS/MS after dansylation. The concentration of CML used is 10 mM, b) Investigation of the stability of CML after coincubation with SpeC for 24 h as measured by HPLC-MS/MS after dansylation. Subscript ds indicates dansylated products, c) MS/MS spectrum of *m/z* 627.2324 at 16.2 min in the chromatogram of the sample taken after 24 h of incubation, d) Kinetics of SpeC-mediated *in vitro* decarboxylation of ornithine as well as e) CML, and f) FmL; n_s_ refers to the number of molecules of substrate. The substrate concentration used is 10 mM. The results are represented as mean and standard deviation of at least three independent replicates.

Instead, the disparity in CML degradation is most probably rooted in their catalytic capacities (Kanjee, Gutsche, Ramachandran, et al., 2011). It has been demonstrated that k*_cat_* of SpeF for the canonical substrate ornithine is more than an order of magnitude lower than that of SpeC (Kanjee, Gutsche, Ramachandran, et al., 2011). Assuming the underground activity of SpeF towards CML undergoes a proportional kinetic reduction, the resulting product formation would fall substantially below the applied analytical detection limits, thereby explaining the apparent lack of activity. This kinetic hypothesis might also explain the observation of marginal CML degradation activity by the SpeF ortholog from *V. cholerae* (Fig. 4a), an organism that naturally lacks a *speC* gene.

The identity of the SpeC reaction products was verified by HPLC-MS/MS as shown in the chromatogram in Fig. 2b, where the peaks corresponding to dansylated CML and CM-Cad are visible, and the mass spectrum of the protonated molecular ion of dansylated CM-Cad in Figs. 2c and S10. We next determined SpeC enzyme kinetics. Using the cognate substrate ornithine, we derived a K_M_ of around 3 mM (Fig. 2d), and calculated a decarboxylation velocity per enzyme of around 6,000 molecules per enzyme and minute both matching previous data (Applebaum et al., 1977; Fonzi & Sypherd, 1985; Kanjee, Gutsche, Ramachandran, et al., 2011). In comparison, with CML we determined a K_M_ comparable to ornithine (∼7 mM) but a turnover rate of only about 4 molecules per enzyme and minute (Fig. 2e). This around 1,000fold lower activity was expected and perfectly aligns with our hypothesis, that CML is decomposed by an underground metabolism (D’Ari & Casadesús, 1998; Rosenberg & Commichau, 2019). Based on our kinetic data we were now also able to estimate the contribution of *E. coli* to CML degradation in the gut. Assuming the bacterium reaches 1% of bacterial cells in the colon (Conway & Cohen, 2015; Sender et al., 2016) with every cell containing up to 103 molecules SpeC (Schmidt et al., 2016), maximally 4% of all CML decarboxylation reactions (based on the minimum daily intake of 10.6 mg CML) (Mark et al., 2014) can be assigned to *E. coli* given a food residence time in the colon of 30 h (see also supplementary material calculation) (Asnicar et al., 2021). Notably, this contribution represents only a fraction of the total degradative potential, as it must be extrapolated to the entire SpeC-harboring microbial community (Fig. S17-S19). Furthermore, the biological efficacy of underground metabolism is driven by chronic substrate availability (Hellwig et al., 2024; Henle, 2005; Khersonsky & Tawfik, 2010). The slow intestinal transit times, coupled with the continuous daily influx of dietary AGEs, allow steady, low-level enzymatic turnover to process significant quantities of CML over time. Crucially, in localized microenvironments or under dysbiotic conditions characterized by Enterobacteriaceae blooms – such as intestinal inflammation (Lupp et al., 2007; Winter & Bäumler, 2014b; Winter et al., 2013) – the functional impact of SpeC-mediated CML degradation is expected to be proportionally amplified.

The chemical differences between CML and ornithine, the cognate substrate of SpeC, hinted to an unexpected high substrate promiscuity and raised the question, which other lysine derivatives can be processed by the enzyme. For further investigation, we chose *N*_ε_-carboxyethyllysine (CEL), which differs from CML only by an additional methyl group and is an equally important AGE (Scheijen et al., 2016). Markedly, the ε-amino group of proteinogenic lysine is not only prone to spontaneous modifications. An arsenal of distinct enzymes targets it in order to regulate protein function by natural genetic code expansion (Lassak et al., 2022). Accordingly, we further selected AcL, FmL, MML, DML and TML (Fig. S6). These naturally occurring modified lysine derivatives are abundant across a wide array of dietary proteins (Jiang et al., 2018; Servillo et al., 2018; Servillo et al., 2014; Walczak et al., 2025), with the epigenetic histone code representing just one prominent example of this structural diversity (Edrissi et al., 2013; Qi et al., 2018). The continuous ingestion of these diverse lysine modifications from whole foods provides a consistent luminal substrate supply. This ubiquitous presence further reinforces the evolutionary selective pressure on the gut microbiota to maintain and utilize specialized degradative pathways. In addition to the listed non-standard amino acids, we also included lysine and arginine as control substrates, both of which were reported to be substrates of SpeC from *Clostridium aceticum* (Tan et al., 2024). While lysine was slowly degraded, arginine is not a substrate of SpeC of *E. coli* (Figs. S6f and S6g). Besides CML, the non-standard amino acids FmL, MML as well as DML were decarboxylated by SpeC, whereas TML, CEL and AcL were not (Figs. S6 and S11). However, kinetic parameters could be derived only for FmL (K_M_ = ∼7.0 mM; k*_cat_* = 5 molecules/minute*enzyme (Figs. 2f, S12)) and are in the same range as for CML. While SpeC exhibits a large promiscuity, the distal region of its active site channel seems to impose distinct structural limits. It tolerates the linear carboxymethyl group of CML as well as small or partially methylated modifications (FmL, MML, DML). By contrast, it sterically excludes bulkier modifications such as the branched carboxyethyl group in CEL or the voluminous trimethylammonium group in TML. Furthermore, the lack of activity towards AcL, in contrast to the smaller FmL, indicates that the additional steric bulk of the acetyl group, potentially combined with the neutralization of the *N*_ε_-charge, might disrupt productive substrate positioning. As discussed earlier, the Fold Type I architecture of SpeC lacks a highly restrictive substrate binding pocket, featuring instead a more extended active site channel (Kanjee, Gutsche, Alexopoulos, et al., 2011; Kanjee, Gutsche, Ramachandran, et al., 2011). To the best of our knowledge, there is currently no crystal structure available that allows for a precise deduction of the spatial and electrostatic constraints within this distal catalytic channel. While available structures of related enzymes offer valuable insights into the general architecture of the catalytic core (Kanjee, Gutsche, Alexopoulos, et al., 2011; Momany et al., 1995), they fail to accurately represent the specific environment during productive substrate binding. This limitation is compounded by the rapid catalytic turnover of canonical substrate (Kanjee, Gutsche, Alexopoulos, et al., 2011; Kanjee, Gutsche, Ramachandran, et al., 2011), which prevents the capture of the pre-catalytic external aldimine state in wild-type enzymes using standard crystallographic techniques (Toney, 2011). However, the newly identified non-standard substrates with their substantially lower turnover rates present a promising tool for future structural studies. Utilizing these slow-reacting derivatives could facilitate the trapping of the enzyme-substrate complex, ultimately allowing for a conclusive mapping of the active site constraints. Beyond these structural considerations, our functional data clearly demonstrate the catalytic breadth of the enzyme. Together, these results establish that SpeC exhibits underground activity toward a variety of non-standard amino acids, thereby generating the previously unknown biogenic amines, formylcadaverine, monomethylcadaverine, and dimethylcadaverine. This in turn raised the question of the selective advantage of degrading such lysine derivatives. We refocused on CML and hypothesized that it might serve as a nutrient source. Our assumption is substantiated by additional metabolites formed during CML utilization (Fig. S13) (Hellwig et al., 2015; Thevaranjan et al., 2017). These observations suggest that CML decomposition follows the same metabolic logic as lysine and ornithine degradation, involving deamination reactions and the subsequent channeling of ammonium into central metabolism (Keseler et al., 2021; Knorr et al., 2018). While the specific enzymes mediating these downstream steps remain to be identified, candidates might include known transaminases involved in polyamine catabolism, such as the putrescine aminotransaminase PatA, or the γ-aminobutyrate aminotransferase PuuE. To discover these molecular players, a comparative transcriptomic approach could identify gene clusters specifically upregulated during growth on CML or CM-Cad. Furthermore, a systematic phenotypic screen of the Keio knockout library (Baba et al., 2006) – mirroring the strategy employed in this study to identify SpeC – followed by *in vitro* validation with purified candidate enzymes, would facilitate a definitive mapping of the complete metabolic pathway.

The physiological operation of this hypothesized catabolic route is directly reflected in the capacity of *E. coli* to utilize CML for its nitrogen requirements (Figs. 3a, S14). Consistently, *E. coli* grew in minimal medium with CML as the sole nitrogen source, showing a marked increase in biomass, evidenced by higher optical density at 600 nm compared to nitrogen-free controls that showed no growth (Fig. 3a, S14). Notably, nitrogen utilization from CML is comparable to lysine (Knorr et al., 2018). Deletion of *speC* resulted in reduced growth. This is plausible, as de-or transamination reactions catalyzed by currently unknown enzymes may also act directly on CML (Hellwig et al., 2019). Nevertheless, the phenotype of the Δ*speC* mutant underlines the importance of the initial decarboxylation step.

**Fig. 3.**
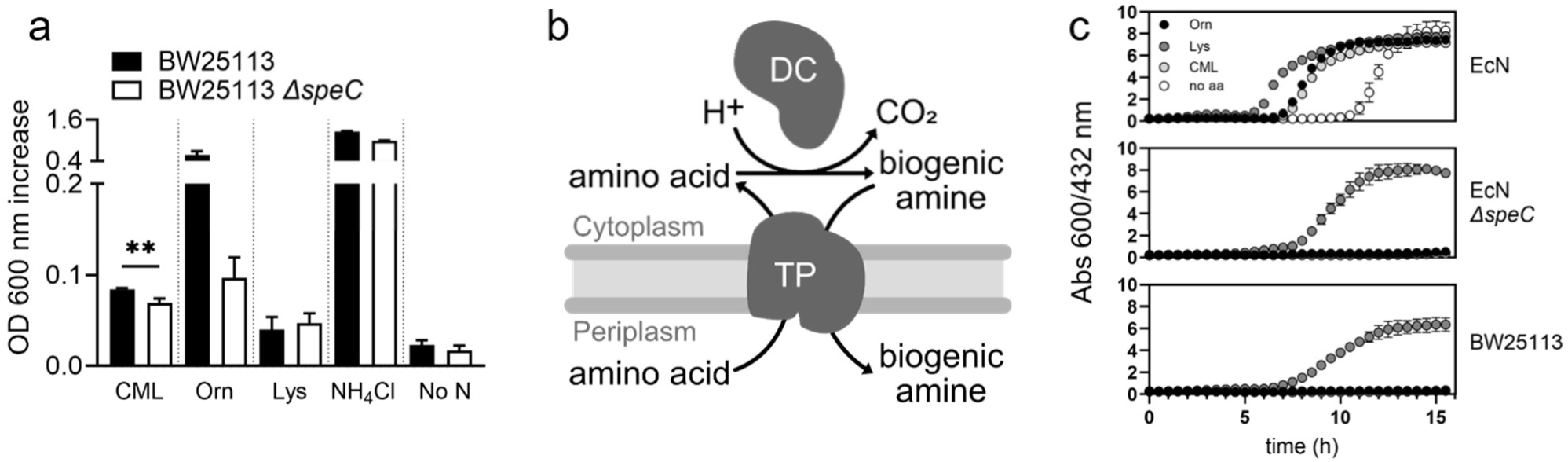
Physiological relevance of CML decarboxylation in *E. coli.* a) Growth analysis of *E. coli* wild type and Δ*speC* mutant cells in M9 minimal media with CML, ornithine, lysine or NH_4_Cl as sole source of nitrogen. Depicted is the maximum increase in OD_600_ over 48 h of incubation at 37 °C. The results were obtained from biological triplicates shown as mean with standard deviation. Unpaired *t*-test was performed with GraphPad Prism 9, yielding a *P-*value of 0.0063. Abbreviations: Orn, ornithine; Lys, lysine; No N, no nitrogen. b) Model of acid stress response mediated by an amino acid decarboxylase and a transporter. The degradation of an amino acid by a decarboxylase (DC) leads to consumption of a proton and production of a more alkaline biogenic amine. This amine is exported by transporter (TP), specifically an antiporter, in turn importing a new amino acid molecule. This process raises the internal and external pH. c) CML-dependent activation of the *E. coli* pH stress response. Investigated was the ability to increase external pH by decarboxylation of ornithine (ODC), lysine (LDC) or CML in *E. coli* Nissle (EcN, ODC^+^/LDC^+^), Nissle Δ*speC*, and BW25113 (ODC^-^/LDC^+^) cells. The restoration of neutral pH was monitored over 16 h through absorbance at 600 and 432 nm of the indicator bromocresol purple that was present in the medium adjusted to pH 5.0. Abbreviations are as a), except for no aa; no amino acid added. The results were obtained from biological triplicates shown as mean with standard deviation

Beyond serving as a nitrogen source, a second possibility is that CML decarboxylation contributes to acid stress resistance in *E. coli* (Jung et al., 2018; Li et al., 2024). Canonically, pH homeostasis is maintained by proton consumption during decarboxylation reactions (Fig. 3b) (Jung et al., 2018). During passage through the gastrointestinal tract, *E. coli* encounters a wide pH range, and depending on the specific niche, different decarboxylase modules are engaged (Schumacher et al., 2023). Whereas the glutamate decarboxylase (Gad) system dominates in the heavily acidic stomach environment, the milder acidity of the colon – where *E. coli* predominantly resides – is counteracted by lysine and ornithine decarboxylation through the CadABC-module or the SpeF/PotE system, respectively (De Biase & Lund, 2015; Kanjee, Gutsche, Ramachandran, et al., 2011). We therefore wondered whether SpeC contributes to this pH-stress response and compared two model strains: K-12, which carries only a functional CadABC-module (https://bacdive.dsmz.de/strain/4414) (Brameyer et al., 2020; Buchner et al., 2015) and the probiotic strain Nissle 1917, which additionally harbors a functional SpeF/PotE system (Sonnenborn & Schulze, 2009). Acid resistance was assessed by classical colorimetric assays using the indicator bromocresol purple (pH transition range 5.2–6.8) (Møller, 1955). As expected, both strains counteracted acid stress in the presence of lysine, whereas only Nissle responded to ornithine (Fig. 3c). Strikingly, CML also proved to be a potent pH-active substrate in Nissle but not in K-12. Unexpectedly, deletion of *speC* abolished pH counteraction not only with CML but also with ornithine, indicating that the previously assumed SpeF/PotE system requires functional SpeC for full activity. Our data further reveals a correlation between the ability to utilize ornithine and the capacity to neutralize acidic pH with CML. Six clinically relevant *E. coli* strains, including enterohemorrhagic and uropathogenic isolates, either did or did not elevate acidic pH with CML depending on their ornithine utilization capability (Fig. S15, Table S2). With SpeF excluded as the CML-decarboxylase, we consider PotE a plausible contributor, either exporting CM-Cad and/or importing CML. In this framework, SpeC provides the catalytic potential, whereas transport and regulation gate whether CML can be used for acid resistance, providing a parsimonious explanation for the strain-specific heterogeneity. Given the substrate promiscuity of SpeC and the likelihood that diet-derived non-standard amino acids reach the colon largely unabsorbed, strains carrying such a module can exploit a general, diet-linked route into acid resistance and nitrogen metabolism.

To ask whether SpeC promiscuity towards CML is *E. coli*-specific (Fig. 4a), we examined orthologs from the key foodborne enterobacterial pathogens *S. enterica* and *Y. enterocolitica* for CML decarboxylation; the panel also included SpeF orthologs as controls (Fig. S16). SpeC orthologs cleaved CML to varying extents, with conversion broadly scaling with phylogenetic proximity to *E. coli* (Fig. 4a). Notably, *V. cholerae* SpeF, which was also included in our study, retained low CML decarboxylation activity. Given that *V. cholerae* naturally lacks a SpeC ortholog, this observation suggests that bacteria possessing only a single ornithine decarboxylase may undergo compensatory adaptations. In such cases, the singular enzyme might face evolutionary pressure to retain or broaden its substrate promiscuity, indicating that noncanonical underground activities can independently arise in distinct decarboxylase lineages. The concept of convergent metabolic flexibility is further corroborated by the obligate anaerobic gut commensal *Cloacibacillus evryensis*, which also degrades CML to identical metabolites like CM-Cad but encodes only distantly related proteins to SpeC (Bui et al., 2019).

**Fig. 4.**
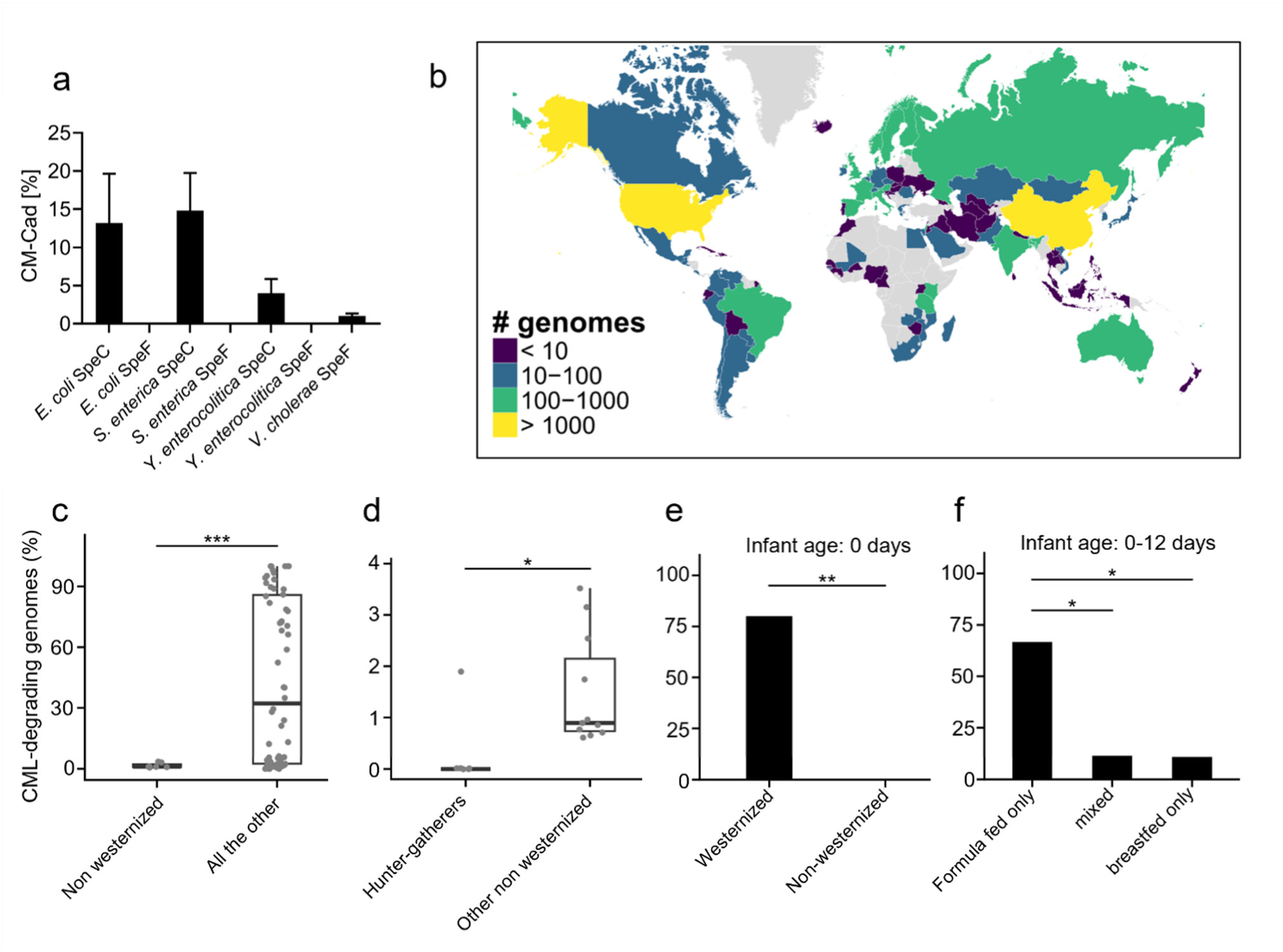
Correlations between the CML degrading microbiota and human diet. a) *In vitro* decarboxylation of CML via SpeC and SpeF orthologs of intestinal pathogens. Note that *V. cholerae* possesses SpeF only. Depicted is the CM-Cad formation relative to the consumed CML after 48 h. The results were obtained from biological triplicates shown as mean with standard deviation. b) Geographic distribution of the number of CML-degrading genomes identified per country/region. A genome deemed to be CML-degrading was defined based on the presence of a SpeC ortholog with a sequence identity ≥70.4% to *E. coli* SpeC. c) Occurrences of CML-degrading bacteria in the human gut microbiota of westernized and non-westernized cohorts. To avoid extreme number and bias, only cohorts with >10 total genomes were included. *P*-value of Student’s t-test is 2.63e^-11^. d) Occurrence of CML-degrading bacteria in the non-westernized cohort, differentiating between hunter gatherer and industrialized community. *P*-value of Student’s t-test is 0.0216. e) Occurrence of CML degrading bacteria in the gut microbiome of infants at 0 days of age, in relationship to the diet of the mother. *P*-value of Fisher’s Exact test is 0.00262. f) Occurrence of CML degrading bacteria in the gut microbiome of infants aged 0 to 12 days based on the type of feeding, differentiating between formula-feeding, breastfeeding or a mixed diet. *P-*values of the Fisher’s Exact test are: 0.0337 (top) and 0.0398 (bottom). The infants of the study in f) are within the westernized group.

Taken together, CML turnover is not confined to *E. coli*: it is widespread across enterobacteria via SpeC orthologs and can be supported at low levels by alternative ornithine decarboxylase architectures.

Guided by these observations, we extended our analysis to the human gut microbiome using the Unified Human Gastrointestinal Genome (UHGG) and Protein (UHGP) collections. To evaluate the distribution of this underground activity across the gut microbiota, we screened for putative CML-degrading SpeC orthologs within the order of Enterobacterales by applying a sequence identity cutoff of 70.4% compared to *E. coli* SpeC. This threshold was intentionally selected as a conservative empirical baseline. It corresponds to the sequence identity of the SpeC ortholog from *Y. enterocolitica*, which was experimentally confirmed to degrade CML in this study (Fig. 4a). The distribution of sequence identities across the analyzed species is visualized in the heatmap in Figure S17. While it is biologically plausible that orthologs with sequence identities below this threshold may also retain CML-degrading activity, restricting the bioinformatic search to this experimentally verified minimum ensures a stringent and robust prediction. Applying this threshold, we identified 10,252 genomes representing 89 species that carry close SpeC orthologs, spanning 29 genera within the Enterobacteriaceae (Figs. S17-S18). Prevalence varied by geography (Fig. 4b): Global mapping revealed that *speC* prevalence is highest in North America, Europe, and Asia, frequently involving continent-specific taxa (Fig. S19). These distribution patterns likely reflect a selective pressure exerted by westernized diets, which are enriched in AGEs such as CML (Fig. 4c). This hypothesis is further supported by the observation that even within non-westernized populations, hunter-gatherer communities – characterized by minimal food processing – exhibit significantly lower prevalence than industrialized non-western cohorts (Fig. 4d). While cooking is a near-universal human practice, it is currently not known how specific culinary techniques lead to different extents of formation of individual AGEs such as CML. Dry-heat methods (e.g. frying or roasting) may generate more and different AGEs than moist-heat preparation (e.g. boiling or steaming). Thus, while non-westernized populations are not AGE-free, their qualitatively distinct exposure likely creates a divergent selective environment for the gut microbiota. Our findings (Fig. 4c,d) might therefore be interpreted as a reflection of these distinct exposure gradients driven by modern food technology (Henle, 2005).

Early-life patterns mirrored these trends: infant microbiomes recapitulated maternal carriage of candidate CML degraders, and formula feeding – relative to mixed or exclusive breastfeeding – was associated with higher prevalence of CML degrading ability in the first days of life (Figs. 4e, f). Collectively, these patterns delineate a widespread, diet-responsive set of SpeC-like AGE decarboxylators in the gut that links processed-food chemistry to microbial programs shaping community structure and function. Motivated by these geography- and early-life gradients – putative ecological readouts of AGE exposure – we next asked whether variation in these taxa also tracks with clinical phenotypes. Across multiple datasets, altered levels of these putative CML-degrading taxa associated with intestinal (e.g. inflammatory bowel disease, colorectal cancer) and hepatic conditions (e.g. fatty liver disease, hepatitis) (Fig. S20). The significant enrichment of SpeC orthologs in these cohorts requires a nuanced interpretation regarding the interplay of diet, microbial metabolism, and disease. We hypothesize that the sustained availability of dietary CML might exert a localized selective pressure, particularly when combined with the physiological alterations of the diseased gut. Intestinal inflammation can cause a significant decrease in luminal pH (Fallingborg et al., 1993; Nugent et al., 2001). Under these acidic conditions, the SpeC-mediated decarboxylation of *N*_ε_-modified lysine derivatives could provide a crucial functional advantage by consuming intracellular protons and supporting pH homeostasis. This metabolic flexibility might aid the survival and characteristic expansion of Enterobacteriaceae frequently observed during intestinal inflammation and advanced liver diseases (Winter & Bäumler, 2014a). Consequently, the elevated prevalence of CML-degrading potential in these cohorts may represent an active metabolic adaptation, driven by the altered gut environment, that enables these bacteria to persist and bloom during host dysbiosis. Although direct *in vivo* validation is required to definitively prove a causal chain, these correlative observations are consistent with the concept that diet-derived Maillard reaction products act as profound ecological drivers of microbiome composition with distinct health implications (Lassak et al., 2023).

Beyond the gut, age-related increases in intestinal permeability elevate circulating CML, which accumulates in microglia and perturbs mitochondrial function (da Cruz Carvalho et al., 2025). CM-Cad is much better internalized by murine microglia (Figs. 5a, b) than CML, indicating that microbiome-derived AGE catabolites can act as brain-facing signals. Whether CM-Cad itself crosses the intestinal barrier is unknown; however, biogenic amines are known to traverse the epithelium by binding to specific receptors on gut epithelial cells (Sudo, 2019), making exposure plausible especially under conditions of increased leakiness with age. Depending on dose and context, CM-Cad could either buffer microglial stress responses or amplify maladaptive activation, outlining a diet-to-microbiome-to-microglia communication axis linking processed-food AGE chemistry to brain innate immunity.

**Fig. 5.**
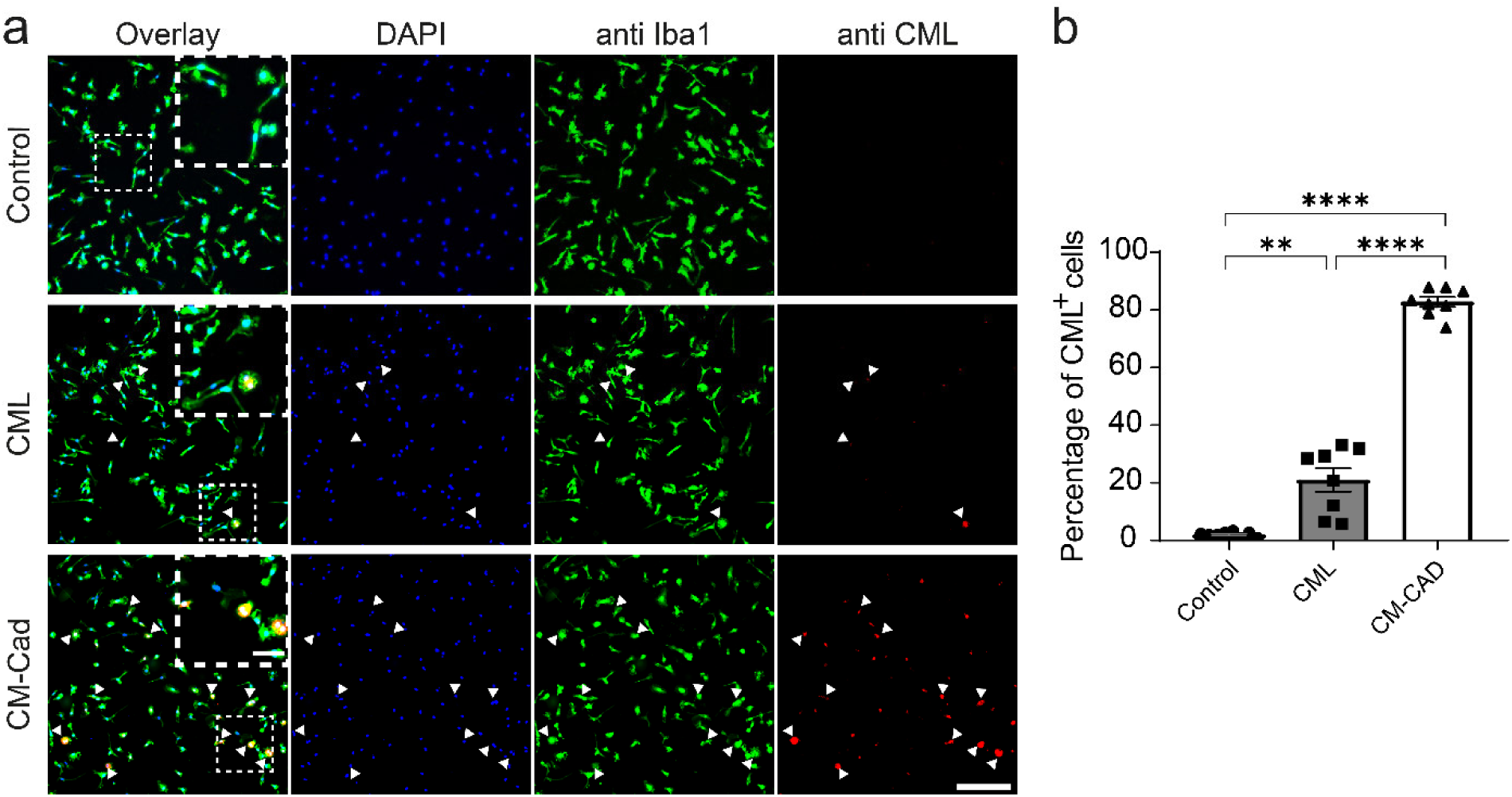
*In vitro* assessment of CML and CM-Cad uptake by murine primary microglial cells isolated from neonatal WT mice (C57BL/6N). The cells were either kept under control conditions or incubated with CML or CM-Cad. a) Representative fluorescent images of murine primary microglia immunolabeled with antibodies against Iba1 (green), CML (red) and CM-Cad (red). DAPI is seen in blue. White arrowheads indicate CML or CM-Cad positive cells. Scale bar 100 µm for the overview and 30 µm for the inset. b) Quantification of the percentage of CML- and CM-Cad positive microglial cells (both detected with an antibody against CML) relative to the total microglial cell number (Iba1^+^). Graph shows mean (±s.e.m.). Each symbol represents data from one experiment with 5×10^4^ cells each, with *n* = 8 biological replicates per experimental group. Significant differences were determined by one-way ANOVA followed by Dunnett’s T3 multiple comparison test, *F* (2.000, 9.534) = 276.5 (\*\**P* = 0.0064; \*\*\*\**P* = <0.0001).

## 4. Conclusion

To conclude, we have identified the enterobacterial SpeC as a promiscuous ornithine decarboxylase that ‘upcycles’ diet-borne *N*_ε_-modified lysine derivatives, including CML, for physiological purposes. While CML can serve as a viable N-source under nutrient-limiting conditions, its physiological significance in the complex environment of the human gut extends beyond basic nitrogen acquisition. In environments where nitrogen is abundant but stress factors such as low pH prevail – particularly during intestinal inflammation – the decarboxylation of CML could provide a crucial mechanism for intracellular pH homeostasis. Thus, this metabolic capacity offers a versatile fitness advantage, functioning as both a nutrient salvage pathway and a critical stress adaptation mechanism that supports commensal persistence and expansion during host dysbiosis. While our study establishes a molecular and biochemical foundation using *E. coli* as a laboratory model, the transition to animal studies will be essential to validate these mechanistic insights *in vivo* and assess the impact of CML-derived metabolites on host physiology. Similarly, while the conservation of SpeC homologs and their distribution patterns point to a novel communication axis between diet, the Enterobacteriaceae in the gut microbiome, and host immunity, these bioinformatic findings remain currently correlative. Future research integrating longitudinal clinical data and functional metagenomics is required to definitively map the causal role of microbial CML metabolism within the human gut. By reframing lysine modifications as actionable inputs for microbial physiology and interkingdom signaling, our work provides a compact framework to interpret – and potentially steer – microbiome-host interactions.

## Acknowledgements

We are grateful to Prof. Dr. Bärbel Stecher for providing the *E. coli* strain UPEC UTI89. We are grateful to Dr. Jinlong Ru for fruitful discussions.

## 5. Statements and declarations

### Author Contributions

The manuscript was written by JL, MH and EFA with contribution from all the authors. The study was designed by JL and MH, with contribution of all the authors. The *in vivo* experiments that led to the discovery of the CML decarboxylase were performed by NG and KIB. The *in vitro* biochemical characterization of *E. coli* SpeC and SpeF was performed by JM, PV and MW. *In vivo* experiments to clarify the physiological relevance of CML degradation in *E. coli* were performed by EFA. The synthesis of the compounds used in this study was performed by MH, and PV. The HPLC-MS measurements and analysis were performed by MH, KIB, MW, and PV. The bioinformatic study on the distribution of CML degrading bacteria in the human gut microbiome was performed by FQ. The *in vitro* experiments using primary murine microglial cells on the uptake of CML and CM-Cad were performed by APVJ and TB. All authors have given approval to the final version of the manuscript.

### Notes

The authors declare no competing financial interest.

## Abbreviations

AGE: advanced glycation end-product
CML: *N*_ε_-carboxymethyllysine
CEL: *N*_ε_-carboxyethyllysine
CM-Cad: carboxymethylcadaverine
CM-APA: carboxymethyl-aminopentanoic acid
MML: *N*_ε_-monomethyllysine
DML: *N*_ε_-dimethyllysine
TML: *N*_ε_-trimethyllysine
FmL: *N*_ε_-monomethyllysine.

## Supplementary material (description)

Table S1 includes the genes annotated as decarboxylases in *E. coli* MG1655. Table S2 includes the phenotype of pH stress response in several *E. coli* strains. Table S3 contains the transitions recorded during MRM measurement of the substrates. Tables S4 to S6 include strains, plasmids and primers used in this study. The full calculation of the contribution of *E. coli* SpeC to CML decarboxylation is provided after the tables. Figures S1-S4 include ^1^H and ^13^C NMR spectra of CML and CM-Cad. Figure S5 contains the HR-MS/MS fragmentation spectra of CM-Cad and CML. Figures S6 and S7 have supporting *in vitro* time series of SpeC in presence of several substrates and time series of SpeF in presence of ornithine respectively. Figure S8 contains the multiple sequence alignment of the ornithine decarboxylases used for *in vitro* tests. Figure S9 contains the comparison of the structures of OdcI and models of SpeC and SpeF. Figures S10 and S11 contain HPLC and MS data of the metabolites incubated in presence of SpeC. Figure S12 includes kinetic curves for SpeC in presence of lysine derivatives. Figure S13 contains the proposed metabolic pathway for CML in *E. coli*. Figures S14 and S15 include supporting growth data to the nitrogen source test and pH stress response curves for clinical *E. coli strains*. Figure S16 includes supporting *in vitro* data proving that SpeC orthologs from other enterobacteria function as ornithine decarboxylases. Figures S17 to S20 include supporting information to the bioinformatic study, including a phylogenetic tree of SpeC orthologs in the human gut microbiota, occurrence of CML degrading bacteria in Enterobacteriaceae, unique species across the different continents and occurrence of CML degrading bacteria in relation to different diseases.

## Supplementary Material

**Table S1.**
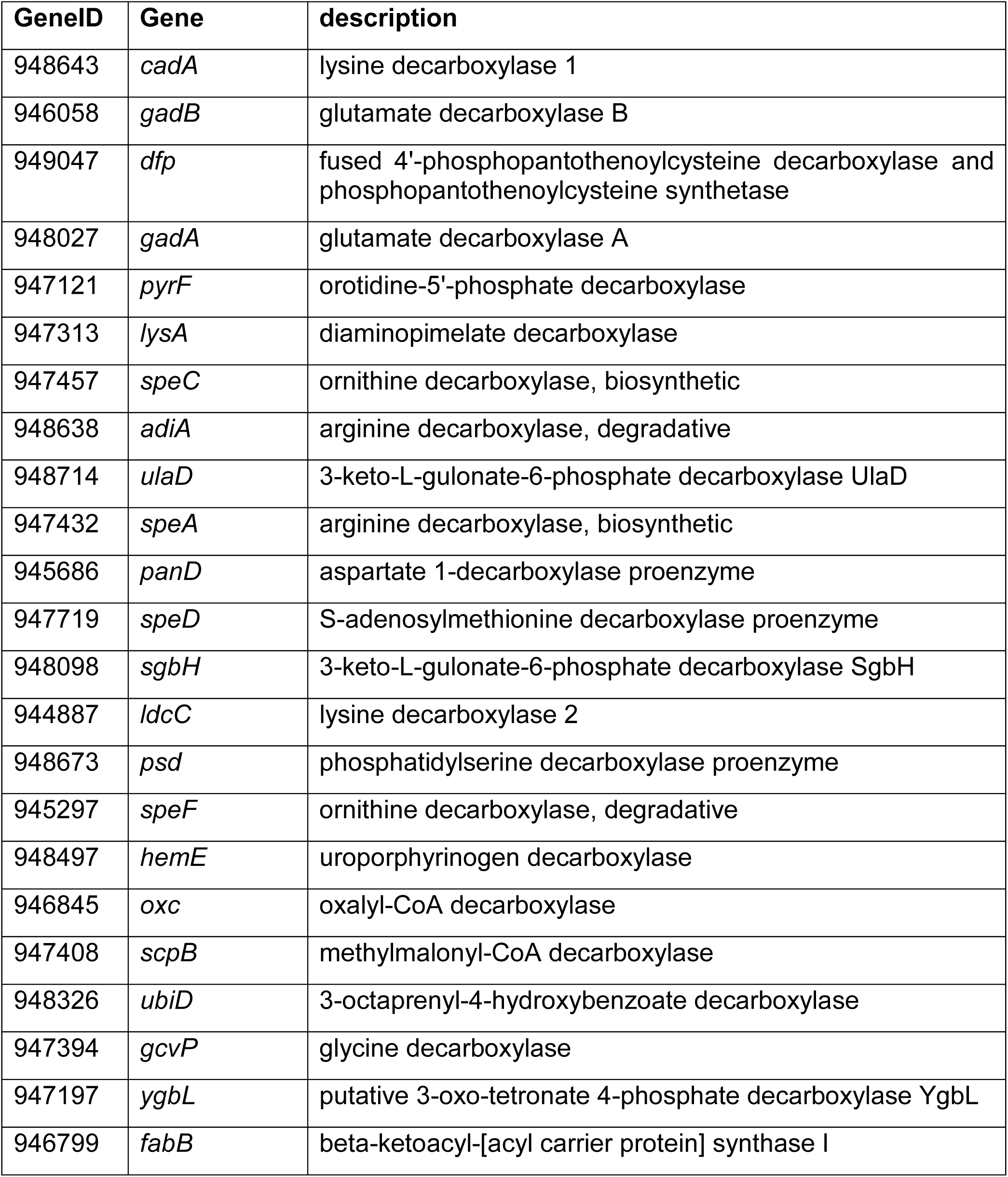
Genes annotated as decarboxylases in *Escherichia coli* MG1655.

**Table S2.** CML dependent pH stress response in the clinical *E. coli* strains, summarized as a table. “++" indicates a saturated bromocresol purple over time and a successful pH stress response, “+” refers to a positive response happening close to the base medium, while "-" refers to absence or strong delay of said curve.

| Strain | Ornithine | CML |
| --- | --- | --- |
| O157:H7 | ++ | ++ |
| ATCC 8739 | + | + |
| O111:H- | + | + |
| UTI 89 | + | + |
| O127:H6 | - | - |
| CFT073 | ++ | ++ |

**Table S3.** Transitions recorded during MRM measurement of CML, CM-Cad, ornithine and putrescine.

|  | Transition | Fragmentor voltage | Collision energy | Q/q |
| --- | --- | --- | --- | --- |
| CML | $m/z$ 205 → 130 | 100 V | 9 eV | Q |
| | $m/z$ 205 → 142 | 100 V | 9 eV | q |
| CM-Cad | $m/z$ 161 → 86 | 100 V | 10 eV | Q |
| | $m/z$ 161 → 98 | 100 V | 10 eV | q |
| Ornithine | $m/z$ 133 → 70 | 100 V | 20 eV | Q |
| | $m/z$ 133 → 116 | 100 V | 5 eV | q |
| Putrescine | $m/z$ 89 → 61 | 100 V | 10 eV | q |
| | $m/z$ 89 → 72 | 100 V | 10 eV | Q |

Q, transition utilized for quantitation (quantifier); q, transition utilized for determination of the presence of the analyte (qualifier).

**Table S4.** Bacterial strains used in this study.

| Strain | Genotype or description | Purpose | Source or reference |
| --- | --- | --- | --- |
| *E. coli* BL21 (DE3) | F<sup>-</sup>, *lon*-11, $\Delta$(*ompT-nfrA*)885, $\Delta$(*galM-ybhJ*)884, λDE3 [*lacI*, *lacUV5-T7 gene 1*, *ind1*, *sam7*, *nin5*], Δ46, [*mal+*]K-12(λS), *hsdS10* | protein purification | (Wood, 1966) |
| *E. coli* DH5α λpir | F<sup>-</sup>, $\Delta$(*argF-lac*)169, ϕ80d*lacZ*58(M15), Δ*phoA*8, *glnX44*(AS), λ-, *deoR*481, *rfbC1*, *gyrA*96 (*NalR*), *recA1*, *endA1*, *thiE1*, *hsdR17*, lysogenized with λpir | plasmid cloning and maintenance | (Taylor, Walker, & McLnnes, 1993) |
| *E. coli* BW25113 | F<sup>-</sup>, $\Delta$(*araD-araB*)567, Δ*lacZ*4787(::rrnB-3), λ-, *rph*-1, $\Delta$(*rhaD-rhaB*)568, *hsdR*514 | *in vivo* study | (Datsenko & Wanner, 2000) |
| *E. coli* JW5482-1 | BW25113; Δ*speC*::*kan* | *in vivo* AGE degradation | KEIO collection (Baba et al., 2006) |
| *E. coli* JW4092-1 | BW25113; Δ*cadA*::*kan* | *in vivo* AGE degradation | KEIO collection (Baba et al., 2006) |
| *E. coli* JW0181-1 | BW25113; Δ*ldcC*::*kan* | *in vivo* AGE degradation | KEIO collection (Baba et al., 2006) |
| *E. coli* JW3485-1 | BW25113; Δ*gadA*::*kan* | *in vivo* AGE degradation | KEIO collection (Baba et al., 2006) |
| *E. coli* JW1488-7 | BW25113; Δ*gadB*::*kan* | *in vivo* AGE degradation | KEIO collection (Baba et al., 2006) |
| *E. coli* JW5731-1 | BW25113; Δ*adiA*::*kan* | *in vivo* AGE degradation | KEIO collection (Baba et al., 2006) |
| *E. coli* JW2905-2 | BW25113; Δ*speA*::*kan* | *in vivo* AGE degradation | KEIO collection (Baba et al., 2006) |
| *E. coli* JW0680-2 | BW25113; Δ*speF*::*kan* | *in vivo* AGE degradation | KEIO collection (Baba et al., 2006) |
| *E. coli* JW2806-1 | BW25113; Δ*lysA*::*kan* | *in vivo* AGE degradation | KEIO collection (Baba et al., 2006) |
| *E. coli* Nissle 1917 | Wild type strain with probiotic properties | Decarboxylase assay | (Hu & Zhang, 2024) |
| *E. coli* Nissle 1917<br>$\Delta speC$ | Nissle, $\Delta speC::kan$ | Decarboxylase assay | This study |
| *E. coli* O157:H7 | Enterohemorrhagic, clinical specimen (human) | Decarboxylase assay | DSMZ No. 17076 |
| *E. coli* O111:H- | Enteropathogenic strain, clinical specimen (human) | Decarboxylase assay | DSMZ No. 8698 |
| *E. coli* ATCC 8739 | Human faecal isolate | Decarboxylase assay | DSMZ No. 1576 |
| *E. coli* O127:H6 E2348/69 | Human infantile diarrhea isolate | Decarboxylase assay | (Iguchi *et al.*, 2009) |
| *E. coli* CFT 073 | Uropathogenic strain, clinical specimen (human) | Decarboxylase assay | (Welch *et al.*, 2002) |
| *E. coli* UTI89 | Uropathogenic strain, clinical specimen (human) | Decarboxylase assay | (Chen *et al.*, 2006) |
| *S. enterica* subsp. *enterica* serovar Typhimurium strain 14028s | Pathogenic bacterium causing gastroenteritis in humans, isolated from a mixture of hearts and liver samples from 4-week old chickens. | genomic DNA extraction for *speC/speF* cloning | (Jarvik, Smillie, Groisman Eduardo, & Ochman, 2010) |
| *Y. enterocolitica* subsp. *enterocolitica* NCTC12982 | Human pathogen isolated from glanders like infection of a 53 years old woman | genomic DNA extraction for *speC/speF* cloning | DSMZ no. 4780 |
| *V. cholerae* O1 biovar El Tor strain N16961 | Human pathogen, agent of cholera. | genomic DNA extraction for *speF* cloning | (Heidelberg *et al.*, 2000) |

**Table S5.** Plasmids used in this study.

| Plasmid | Purpose | Selective marker | Reference |
| --- | --- | --- | --- |
| pBAD33 | Bacterial expression vector; arabinose inducible araBAD promoter (PBAD) | Chloramphenicol resistance | (Guzman, Belin, Carson, & Beckwith, 1995) |
| pET-SUMO | Expression of *E. coli speC* | Kanamycin resistance | Thermo Fisher Scientific |
| pBAD33-*speC* | Expression of *E. coli speC* | Chloramphenicol resistance | this study |
| pBAD33-*speC*<br>*Y. enterocolitica* | Expression of *Y. enterocolitica speC* | Chloramphenicol resistance | this study |
| pBAD33-*speF*<br>*Y. enterocolitica* | Expression of *Y. enterocolitica speF* | Chloramphenicol resistance | this study |
| pBAD33-*speF*<br>*V. cholerae* | Expression of *V. cholerae speF* | Chloramphenicol resistance | this study |
| pBAD33-*speC*<br>*S. enterica* | Expression of *S. enterica speC* | Chloramphenicol resistance | this study |
| pBAD33-*speF*<br>*S. enterica* | Expression of *S. enterica speF* | Chloramphenicol resistance | this study |
| pKD13 | Template for generation of a kanamycin resistance cassette | Kanamycin resistance | Baba *et al.* 2006 (Baba *et al.*, 2006) |
| pRed/ET (pKD46) | Recombinase bearing plasmid for insertion of a kanamycin resistance cassette | Ampicillin resistance | Quick and Easy *E. coli* Gene Deletion Kit, Gene Bridges |
| pET-SUMO-*speC* | Protein purification of *E. coli SpeC* | Kanamycin resistance | this study |
| pET-SUMO-*speF* | Protein purificaion of *E. coli SpeF* | Kanamycin resistance | this study |

**Table S6.**
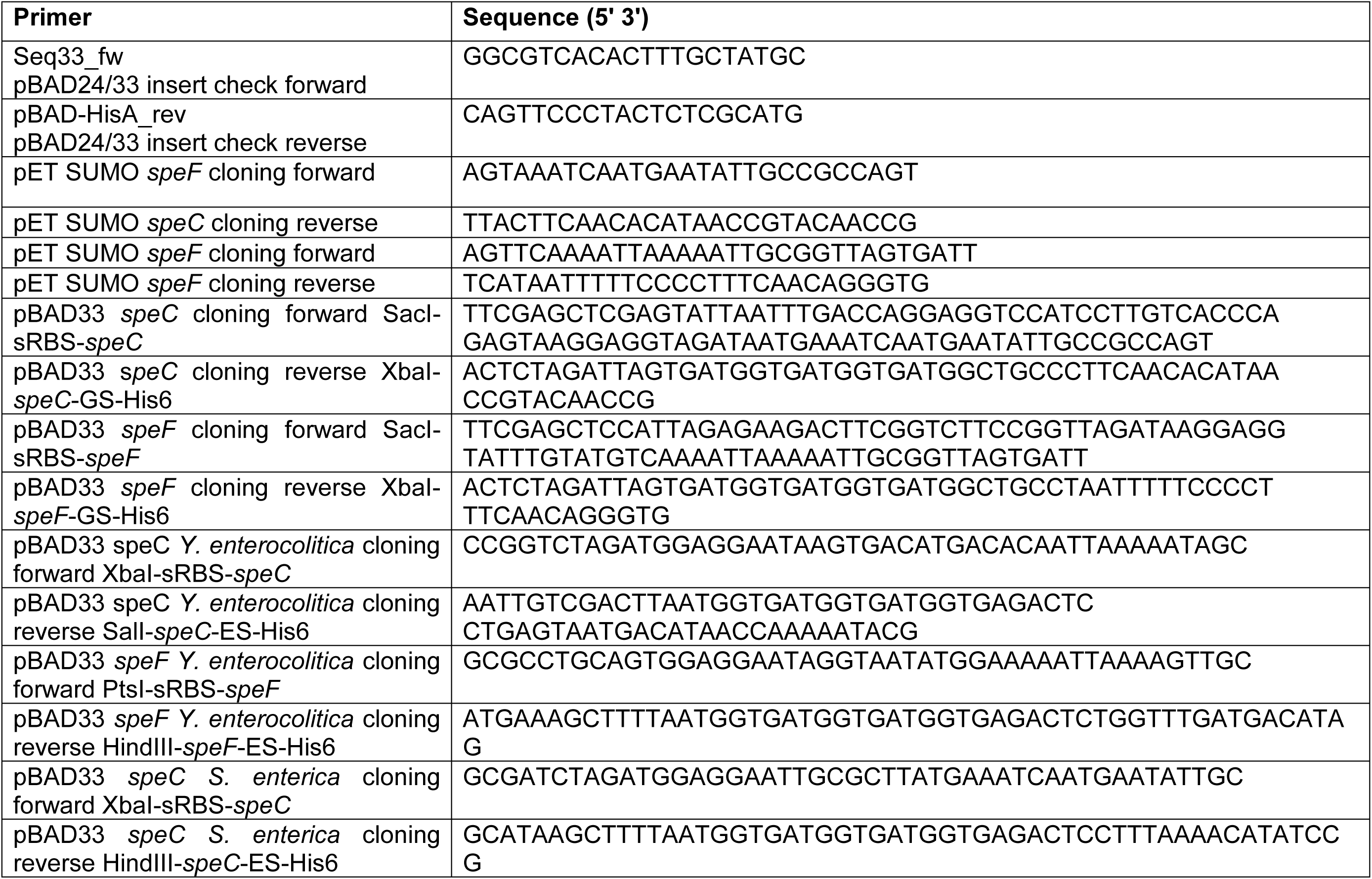
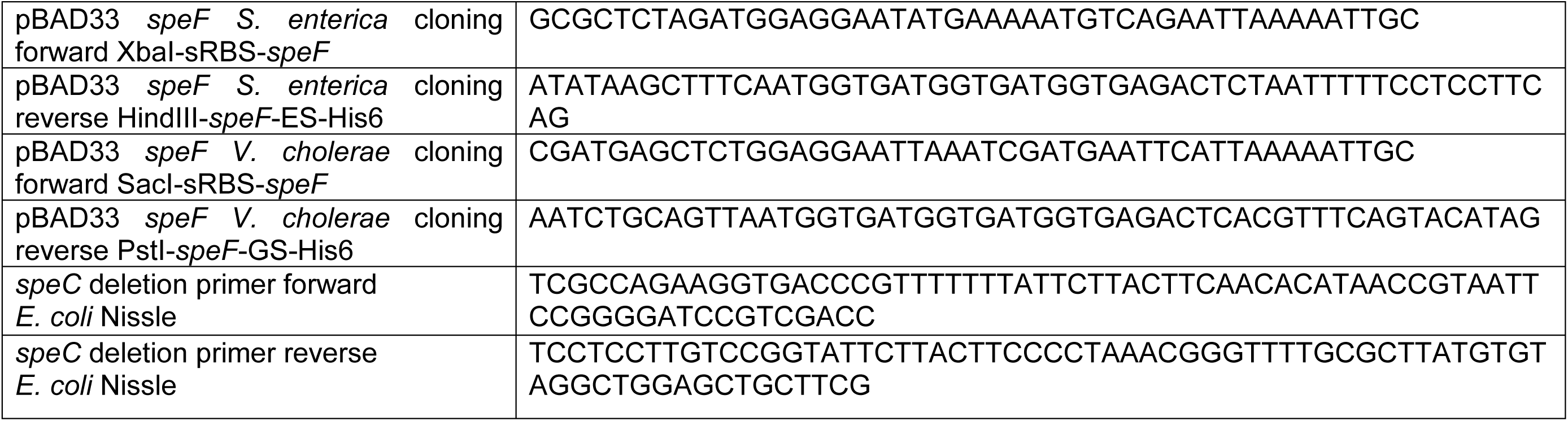
Primers used in this study.

### Calculation of the contribution of *E. coli* SpeC to the decarboxylation of CML in the gut

To obtain a robust estimate for the absolute number of *E. coli* cells in the human colon, we integrated two distinct calculation approaches based on data for a 70 kg reference man (Conway & Cohen, 2015; Sender, Fuchs, & Milo, 2016). Both methods assume a relative abundance of 1% *E. coli* within the total gut microbiota. The first approach is based directly on the total bacterial cell count of 3.8 x 10^12^. Applying the 1% abundance ratio results in a population of 3.8 x 10^11^ *E. coli* cells. The second approach utilizes the total bacterial mass, which is estimated at approximately 0.2 kg. A 1% share of this mass corresponds to 2 g of *E. coli*. Using the globally recognized standard reference mass of 1 pg (10^-12^ g) per *E. coli* cell (bionumbers ID: 101789), this mass-based calculation yields 2.0 x 10^12^ cells. To minimize methodological bias and account for variations in cell size and density, we used the arithmetic mean of these two estimates. This results in a final value of 1.19 x 10^12^ *E. coli* cells, which was used for all subsequent calculations in this study. Based on this value, the metabolic capacity for the degradation of CML by the bacterial enzyme SpeC was calculated. Assuming an enzyme concentration of 103 SpeC copies per cell (Schmidt et al., 2016) and a turnover rate of about 4 molecules per minute, a passage time of 30 hours (Asnicar et al., 2021) yields a theoretical turnover of 8.83 x 10^17^ CML molecules. This corresponds to a mass of approximately 0.3 mg of CML. The microbial degradation potential was evaluated against a reference value of 7.49 mg CML. This value represents the fraction of CML entering the metabolic pathway, derived from a minimal daily dietary intake scenario assuming a 70% metabolization rate and 30% excretion (Mark et al., 2014). Based on these parameters, the theoretical contribution of E. coli to total CML metabolism is approximately 4.0%.

**Figure S1.**
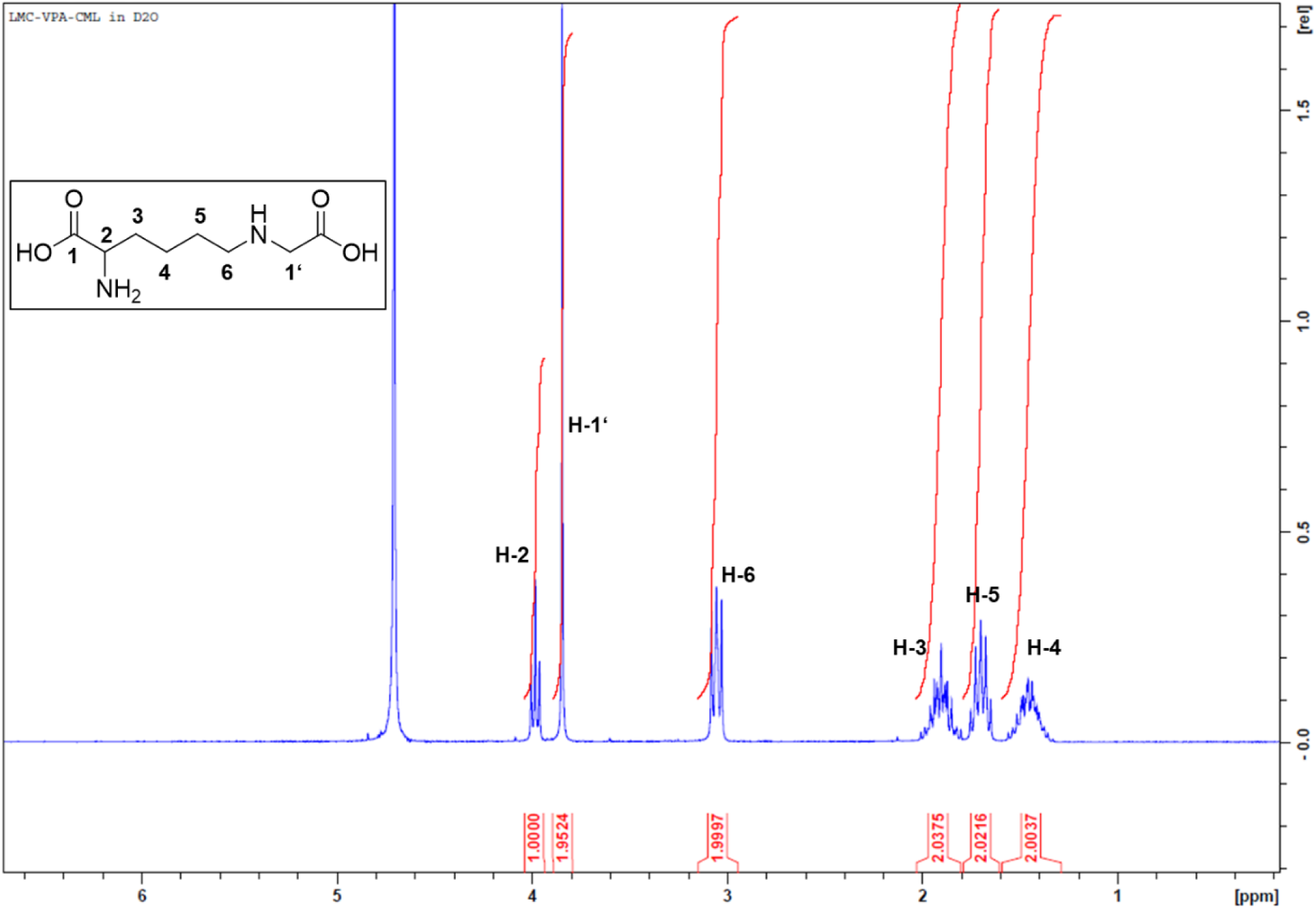
^1^H NMR spectrum of CML.

**Figure S2.**
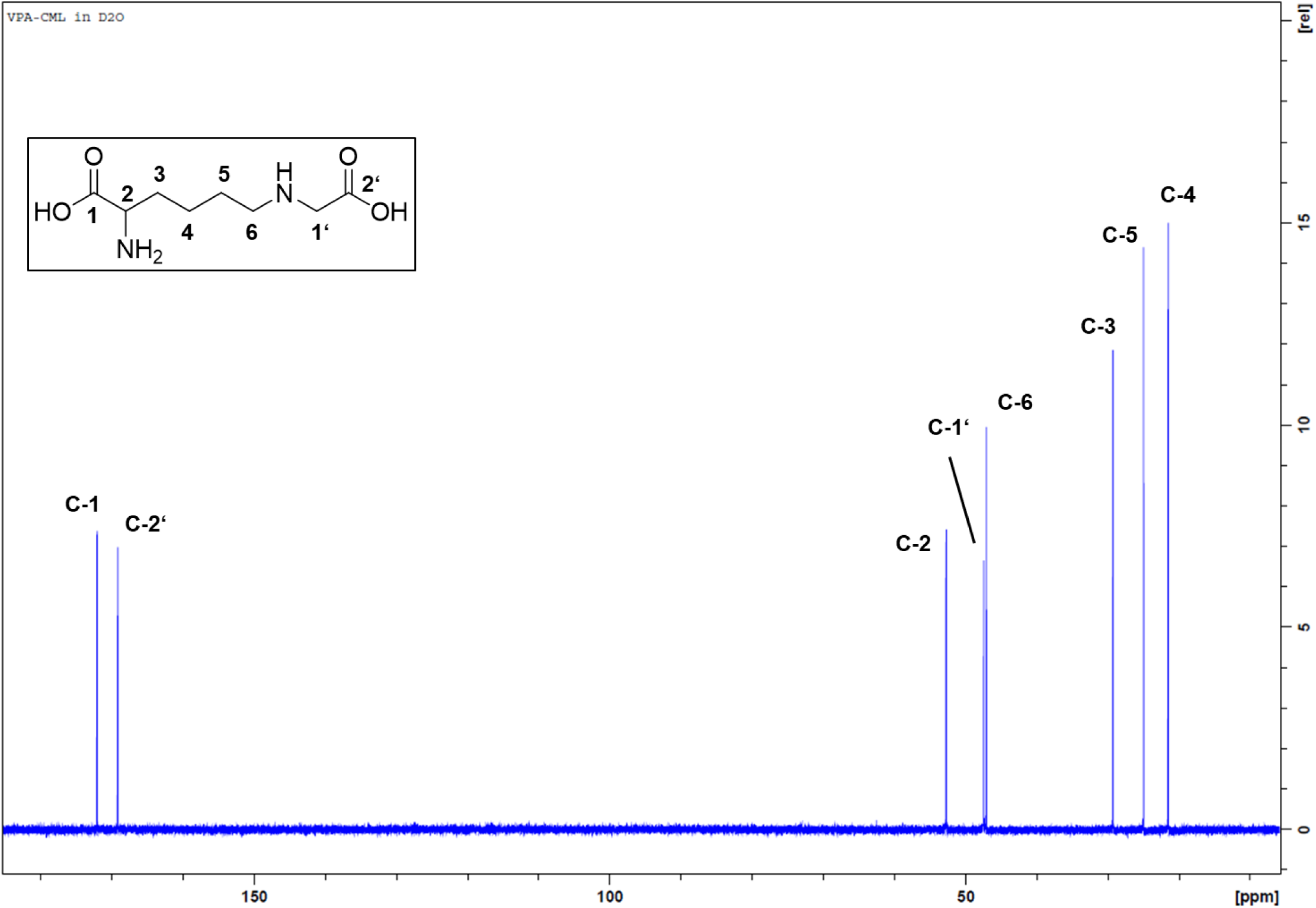
^13^C NMR spectrum of CML.

**Figure S3.**
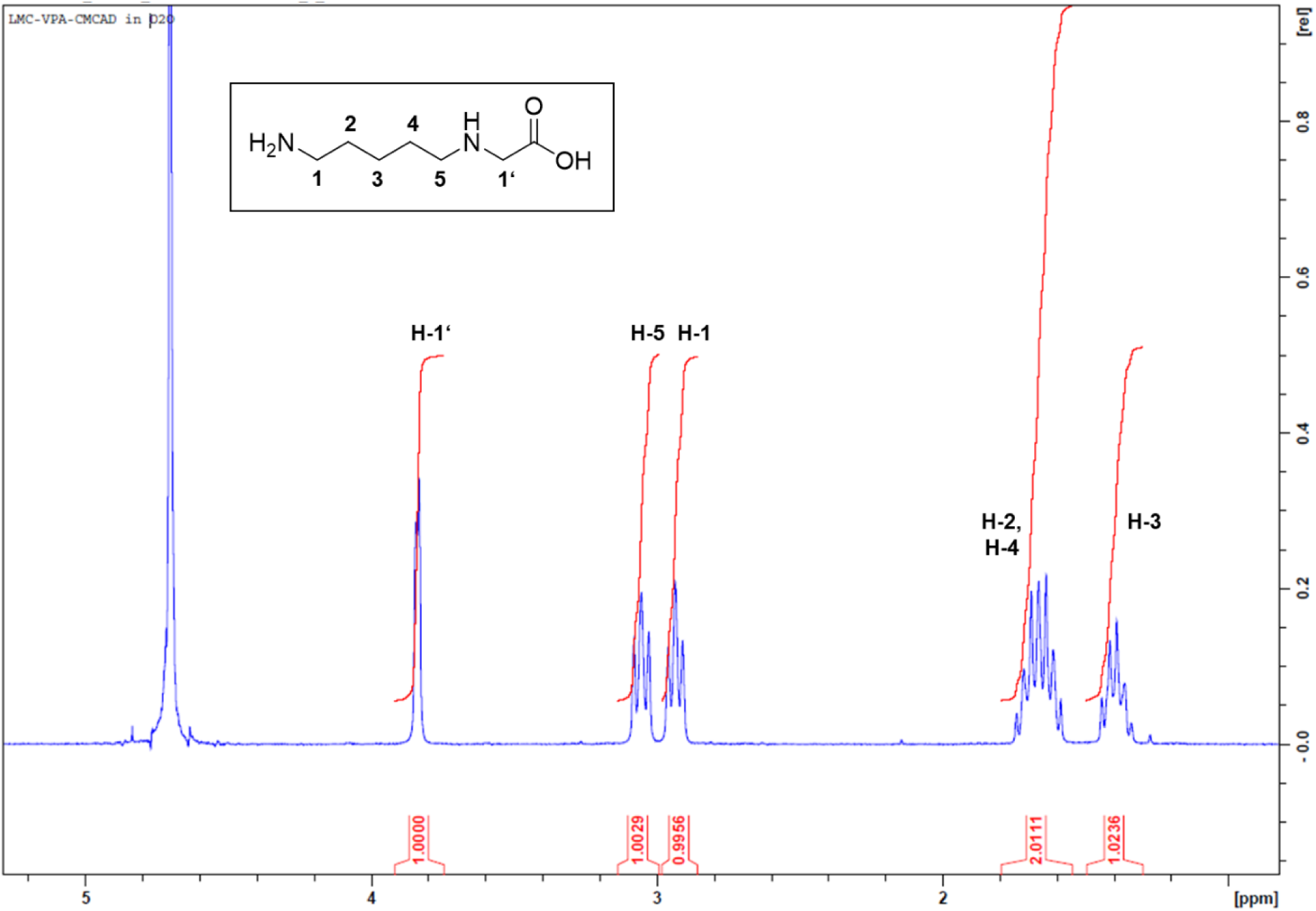
^1^H NMR spectrum of CM-Cad.

**Figure S4.**
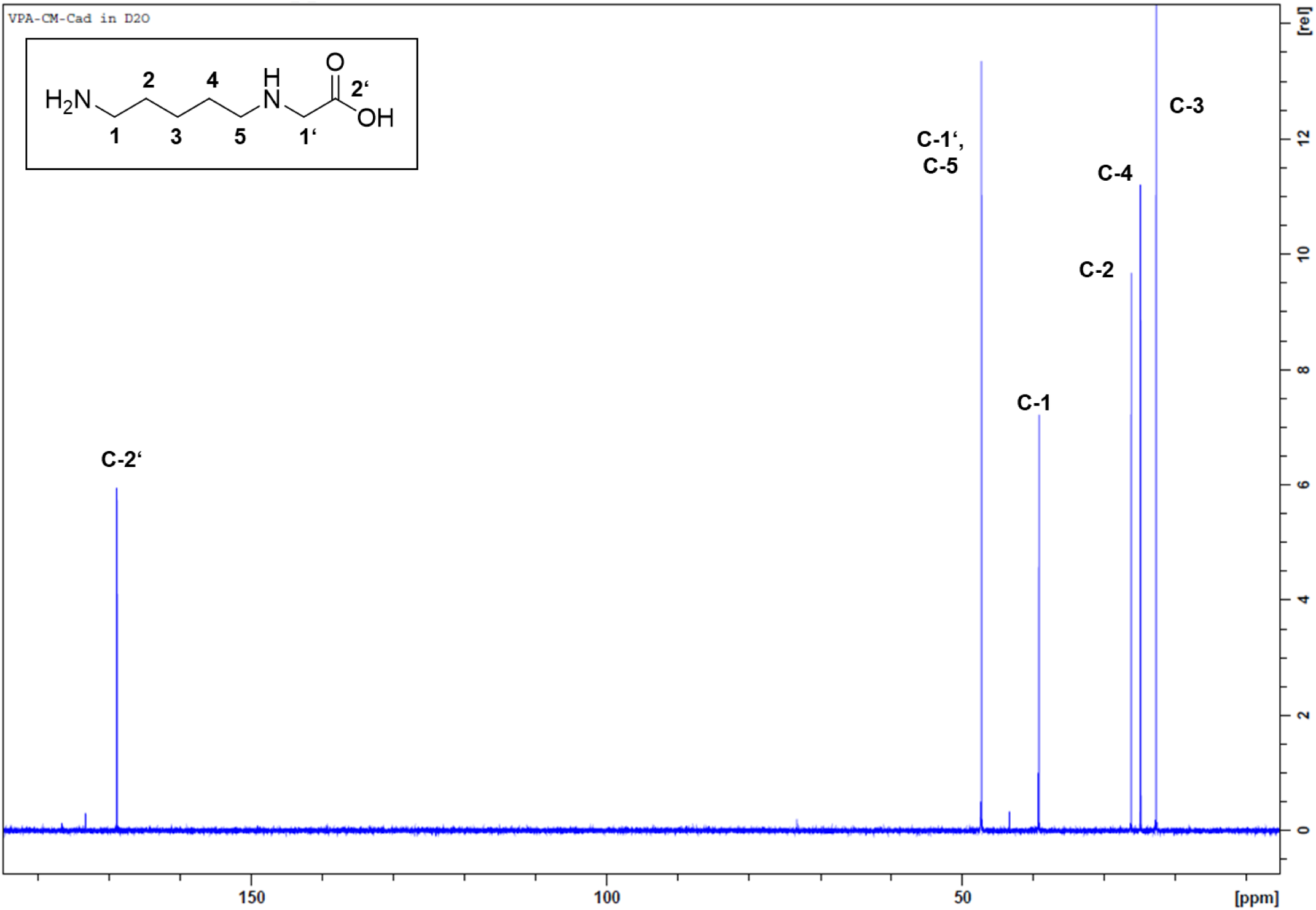
^13^C NMR spectrum of CM-Cad.

**Figure S5.**
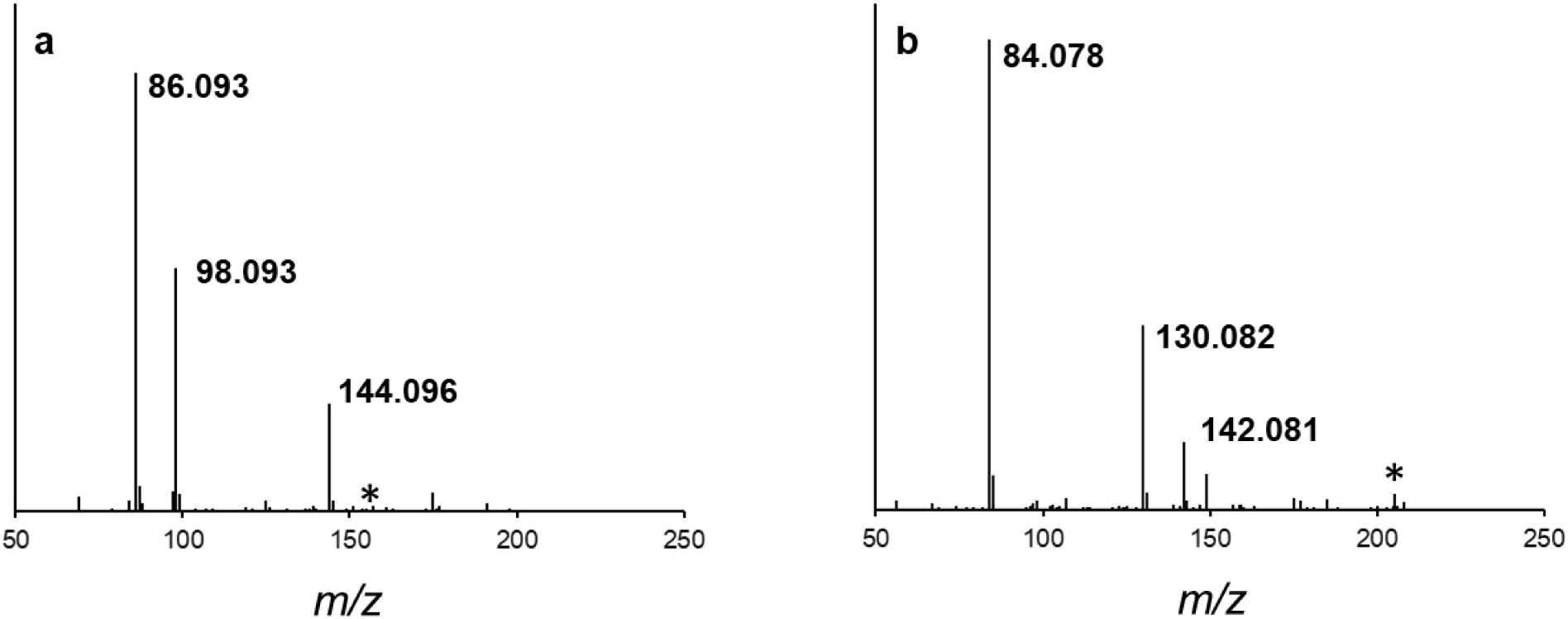
HR-MS/MS fragmentation spectra (80 V, 20-50 eV) of (a) CM-Cad, and (b) CML. Position of the fragmented ion is marked by an asterisk.

**Figure S6.**
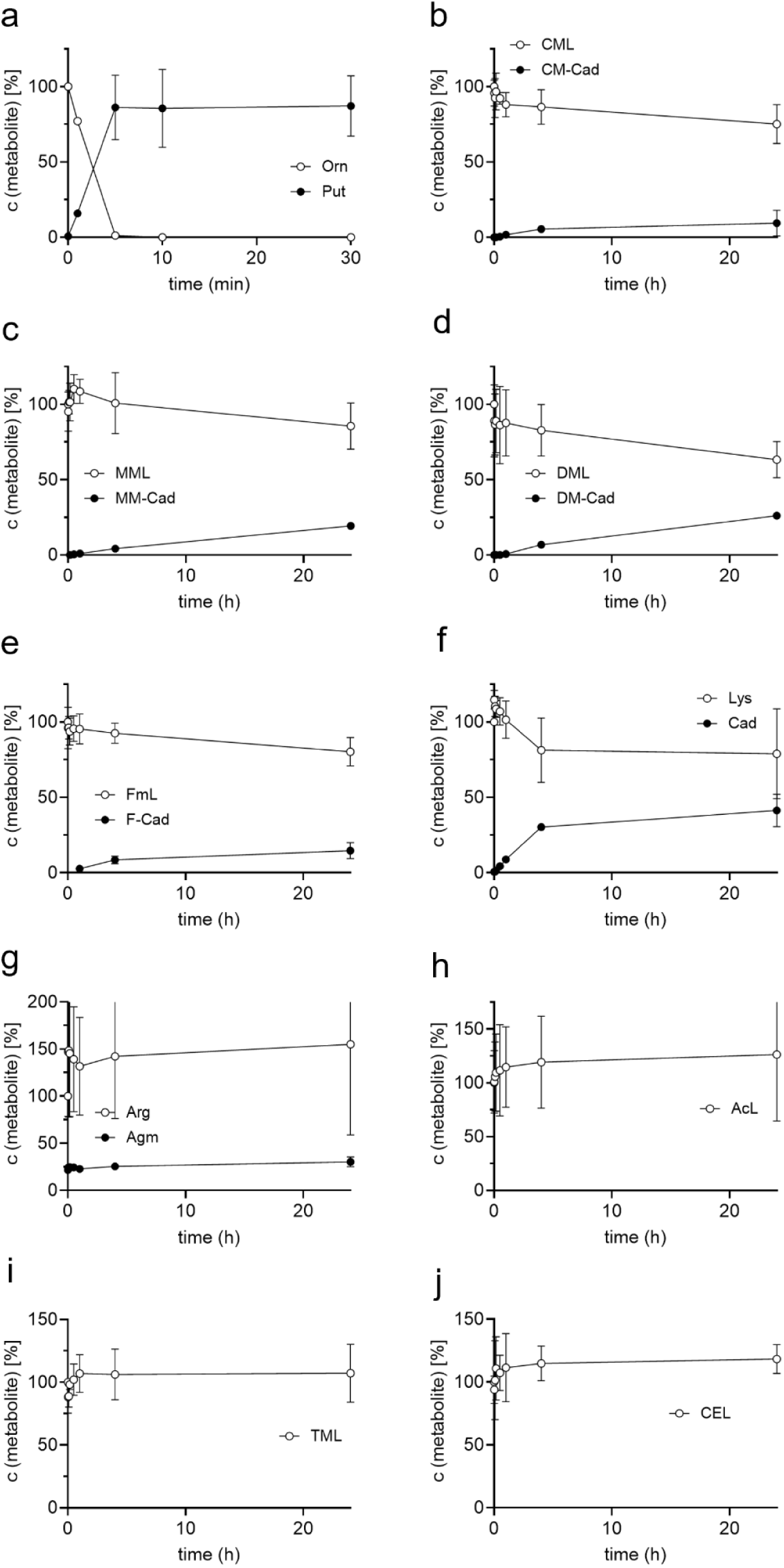
Time course of *in vitro* reactions of purified *E. coli* SpeC with different substrates. Relative amount of (a) ornithine, (b) CML, (c) MML, (d) DML, (e) FmL, (f) L-lysine, (g) L-arginine, (h) AcL, (i) TML, (j) CEL. The metabolites were measured by HPLC-UV after pre-column dansylation, and relative amounts of substrates and their degradation products are shown over time. The enzyme was applied in a concentration of 50 nM for ornithine and 500 nM for the other substrates. The results are represented as mean and standard deviation of three independent replicates.

**Figure S7.**
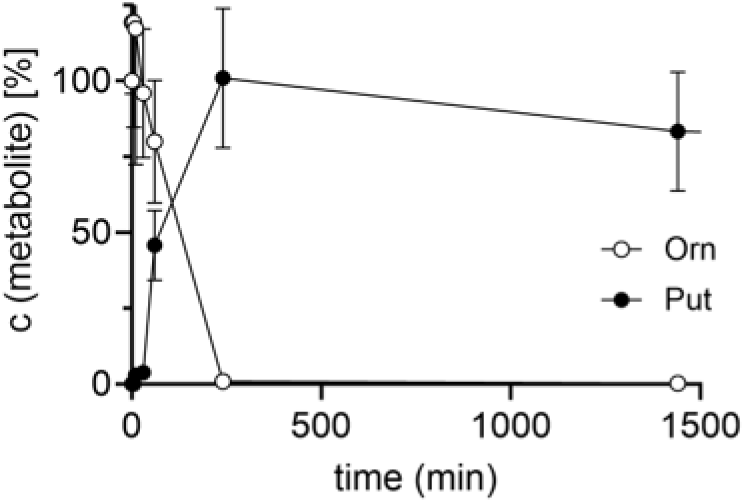
Time course of *in vitro* reaction of purified *E. coli* SpeF in presence of 10 mM ornithine. The metabolites were measured by HPLC-UV after pre-column dansylation. The relative amount of substrate and product is shown. The enzyme was applied in a final concentration of 50 nM. Orn, ornithine; Put, putrescine. The results are represented as mean and standard deviation of three independent replicates.

**Figure S8.**
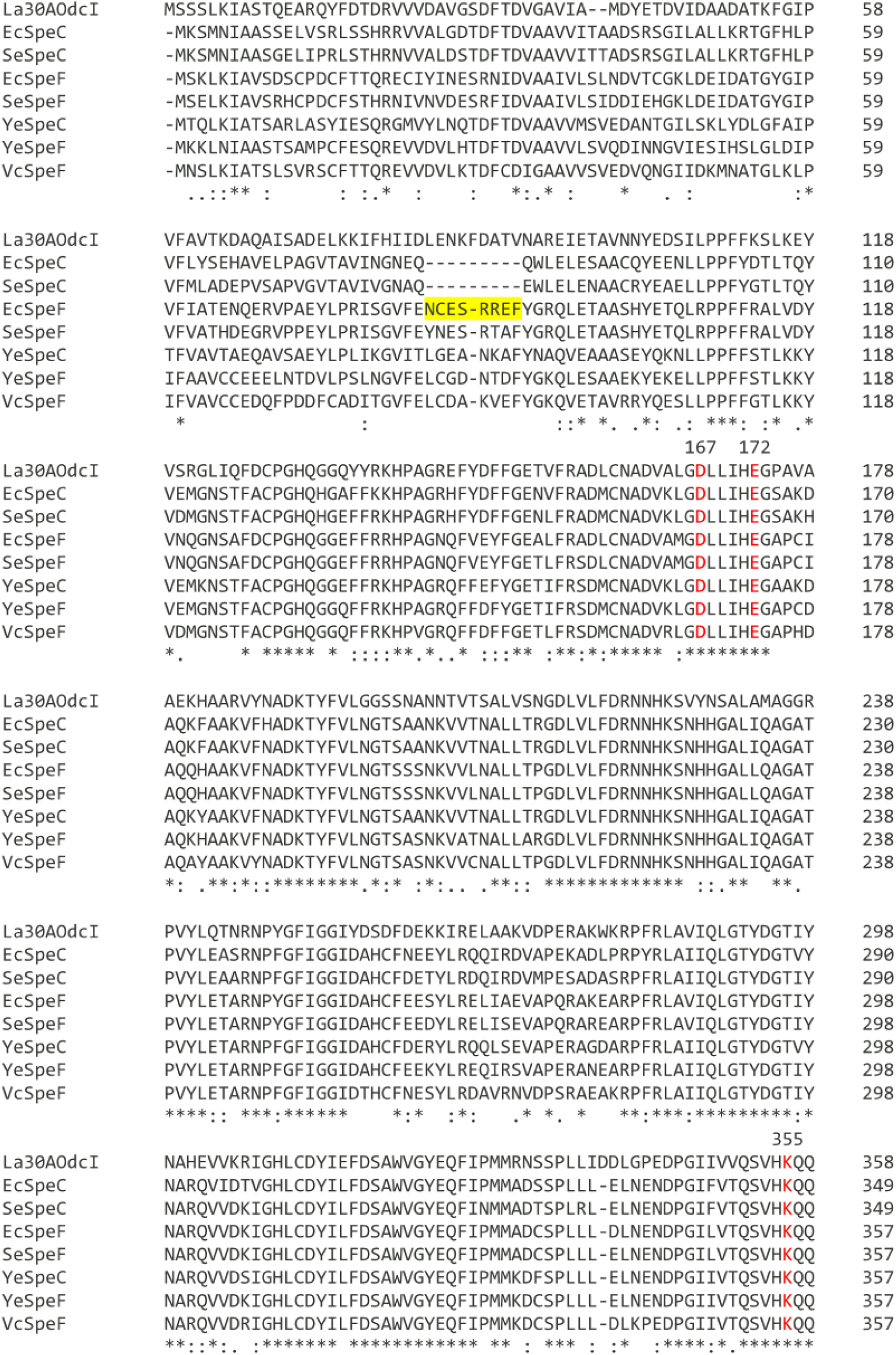
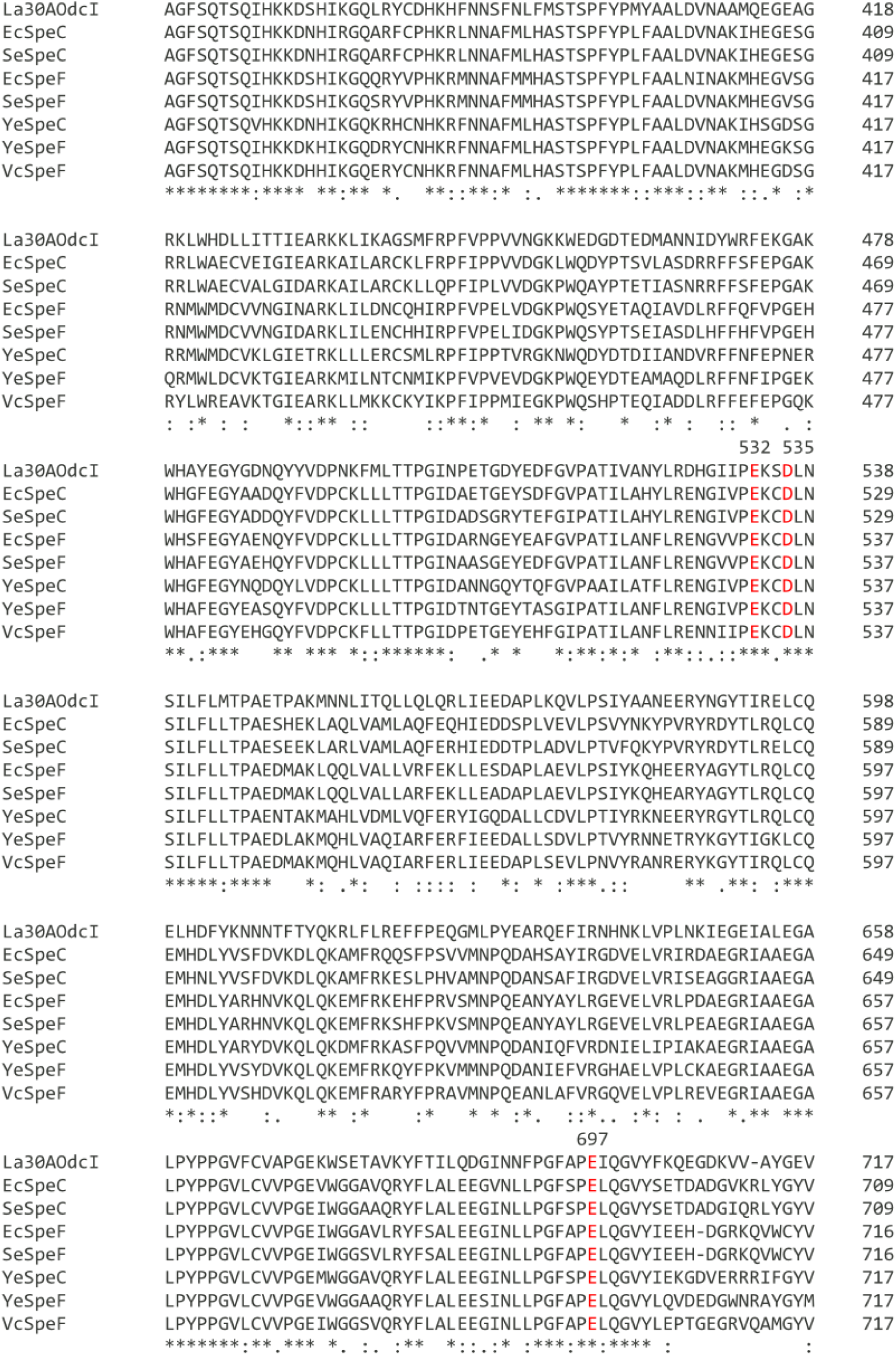
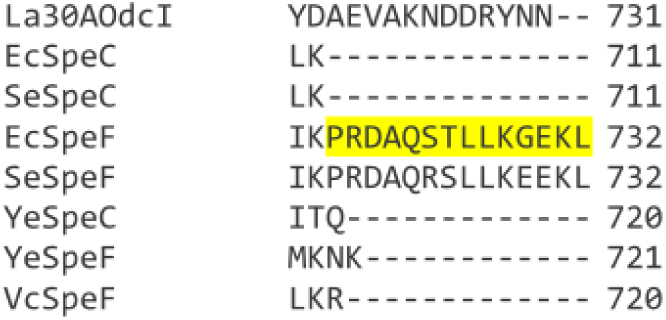
Multiple sequence alignment of the ornithine decarboxylases used for the *in vitro* degradation of CML. OdcI from *Lactobacillus* 30A (UniProt ID P43099) was included to compare key residues identified in previous work (Kanjee et al., 2011). Putative substrate-binding residues are marked in red, comprising the PLP-binding lysine (K355) and negatively charged residues within the substrate-binding pocket (D167, E172, E532, E535, and E697). The alignment was generated using Clustal Omega. Standard symbols indicate conservation levels: fully conserved (*); strongly conserved, scoring > 0.5 in the Gonnet PAM 250 matrix (:); weakly conserved, scoring ≤ 0.5 and > 0 (.); and not conserved (unmarked). Red positions are numbered according to the *Lactobacillus* 30A OdcI sequence. Sequence stretches present in *E. coli* SpeF but absent in SpeC are highlighted in yellow. Abbreviations: *Lactobacillus* 30A, La30A; *Escherichia coli*, Ec; *Salmonella enterica*, Se; *Yersinia enterocolitica*, Ye; *Vibrio cholerae*, Vc.

**Figure S9.**
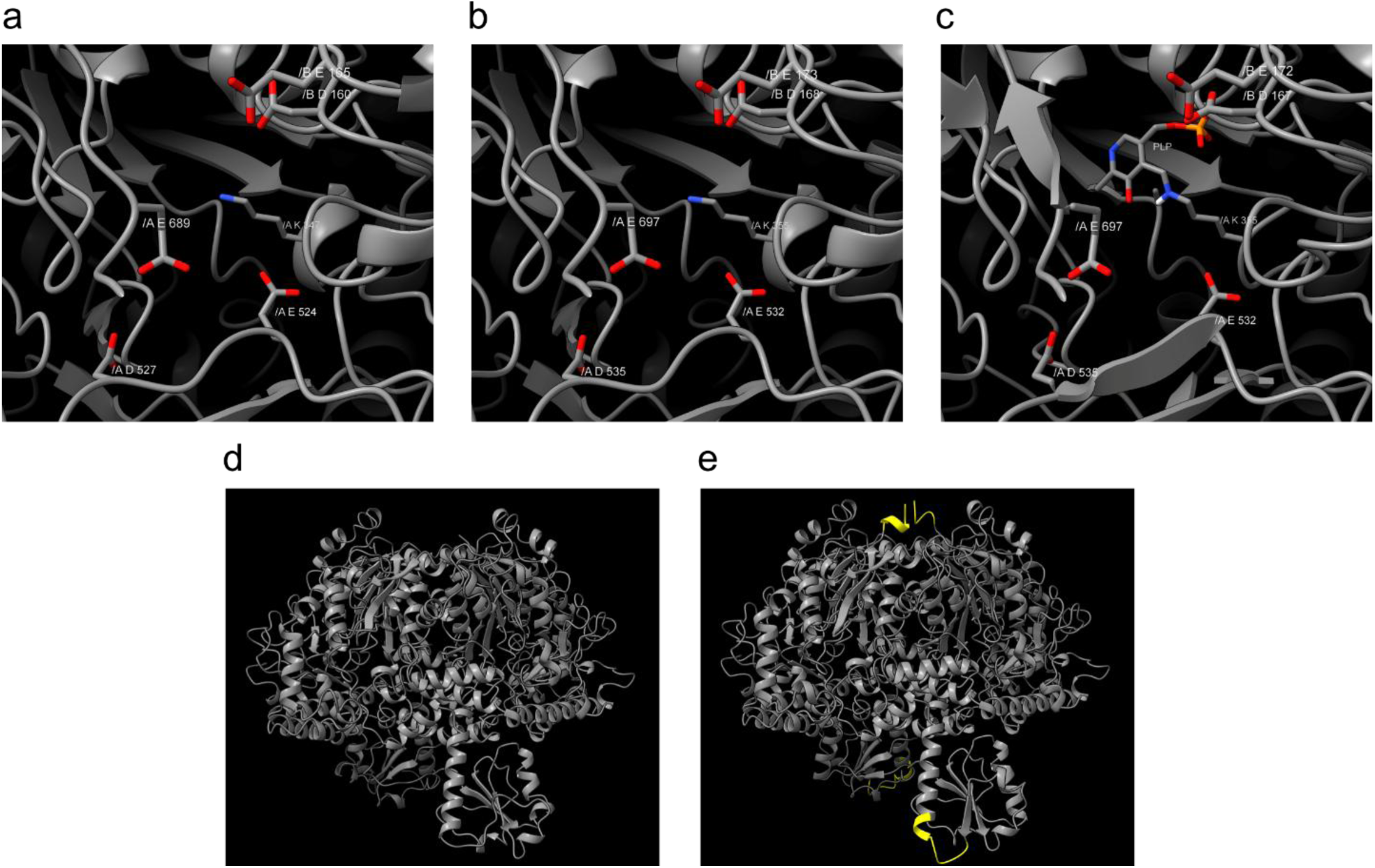
Comparison of the structural models of a) SpeC (uniprot ID P21169, SWISS-MODEL P21169_2-711:1ord.1.B) and b) SpeF (uniprot ID P24169, SWISS-MODEL P24169_2-723:1ord.1.B) from *E. coli* with the crystal structure of c) Lactobacillus 30A OdcI (Uniprot ID P43099, PDB:1ORD). In the first row the PLP binding pocket is showcased with the PLP-binding lysine and the negatively charged residues, for a) SpeC, K347, D160, E165, E524, D527, E689; b) SpeF, K355, D168, E532, D535, E697 and c) OdcI from *Lactobacillus* 30A, PLP, K355, D167, E172, E532, D535, E697. The chain of each residue is specified as A or B. d) Model of SpeC compared to e) SpeF, in which the regions marked in yellow are absent from SpeC, as labeled in the multiple sequence alignment in figure S3. The figure was created using ChimeraX.

**Figure S10.**
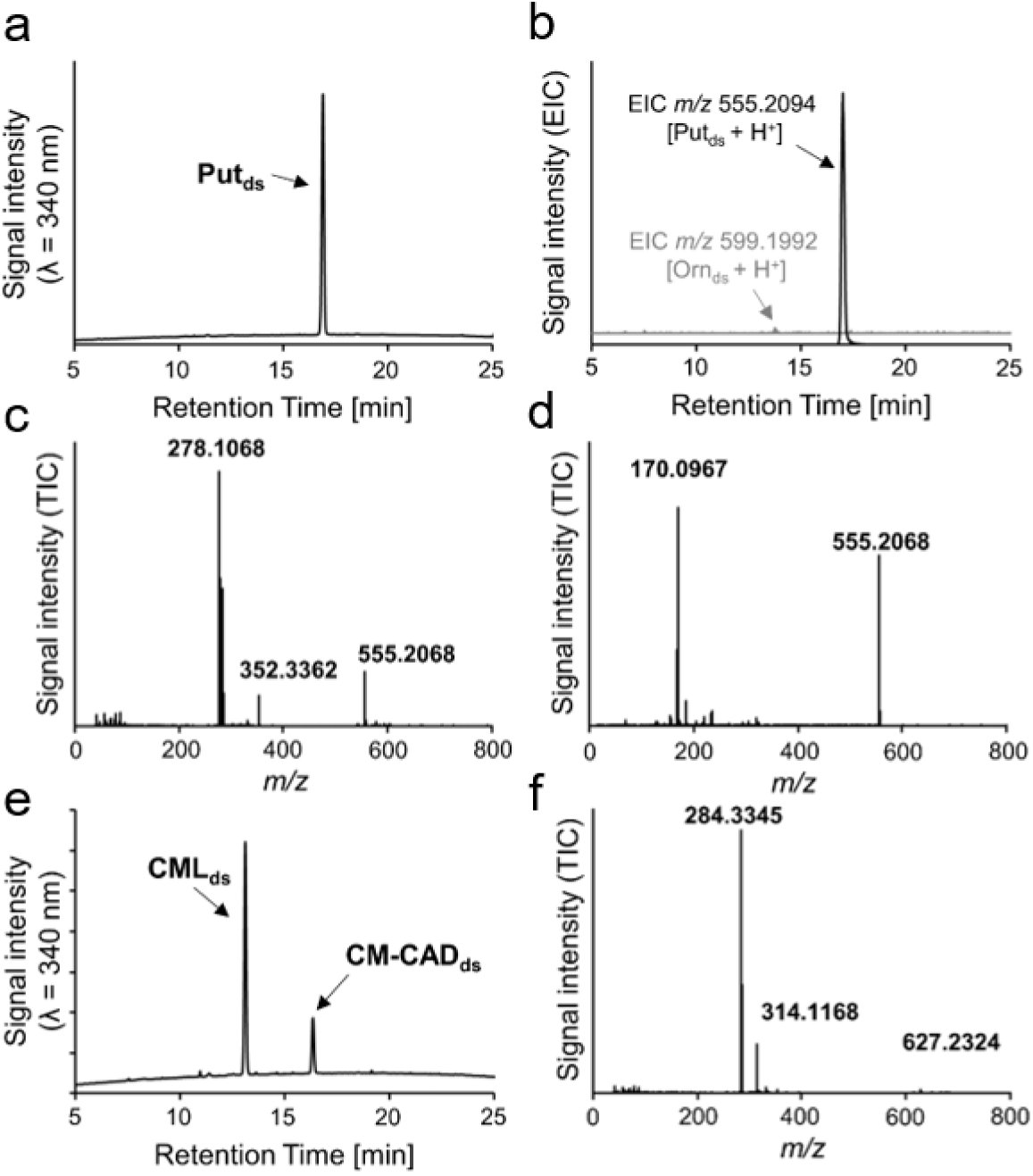
Investigation of the stability of ornithine after coincubation with SpeC for 24 h as measured by (a) RP-HPLC with UV detection and (b) MS/MS detection after dansylation. Please note that the chromatogram of EIC *m/z* 599.1992 is ten times magnified as compared to that of *m/z* 555.2094 in order to see the peak of Orn_ds_ c) Mass spectrum of the peak eluting after 16.9 min in the chromatogram of the sample taken after 24 h of incubation. (d) MS/MS spectrum of *m/z* 555.2068 at 16.9 min in the chromatogram of the sample taken after 24 h of incubation. Abbreviations: Put, putrescine; Orn, ornithine; Subscript ds indicates dansylated product. Investigation of the stability of CML after coincubation with SpeC for 24h as shown in (e) RP-HPLC with UV detection and (f) Mass spectrum of the peak eluting after 16.2 min in the chromatogram of the sample taken after 24 h of incubation.

**Figure S11.**
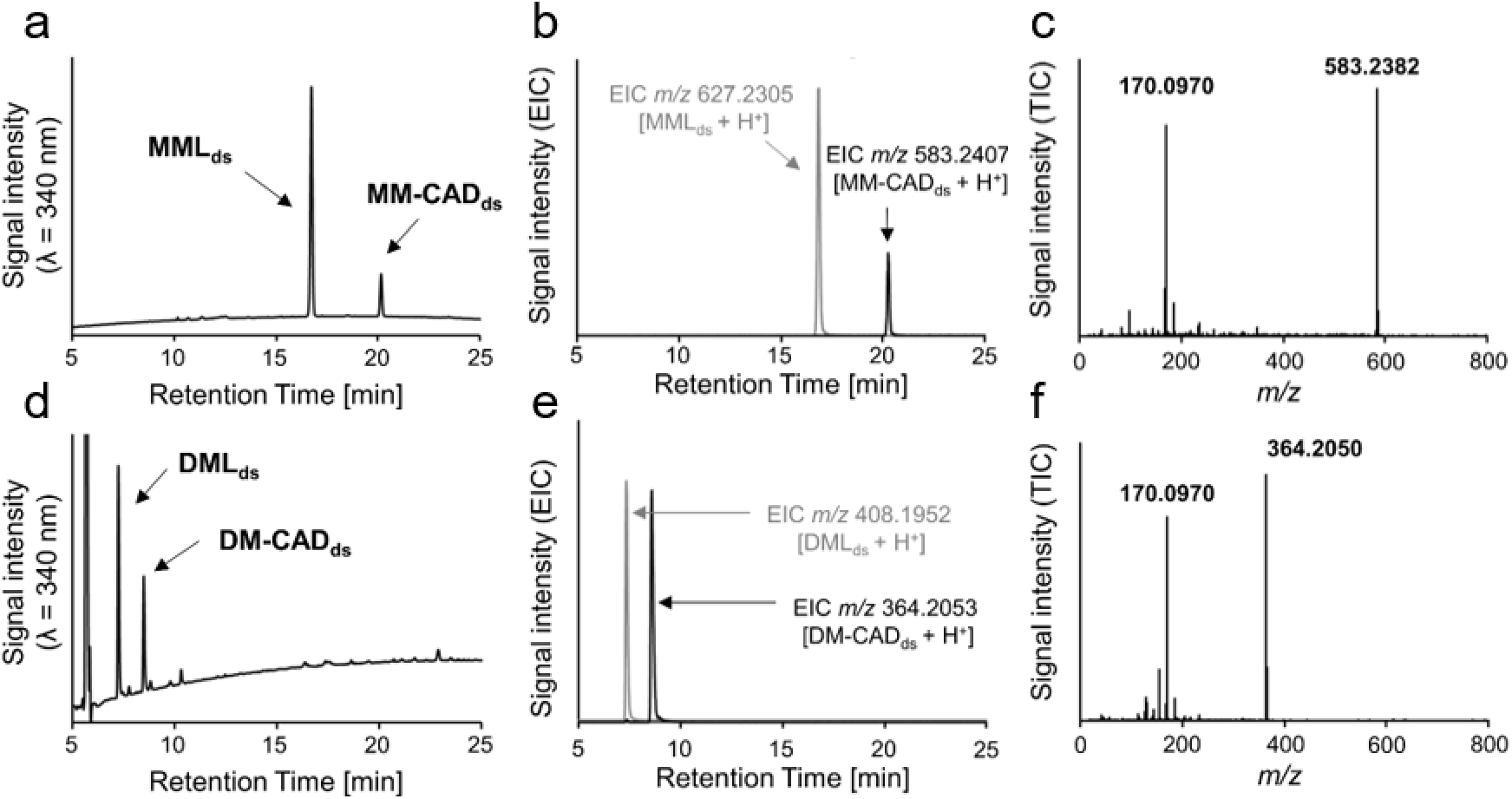
Investigation of the stability of MML and DML after coincubation with SpeC for 24 h. For MML: (a) RP-HPLC with UV detection and (b) MS/MS detection after dansylation. (c) MS/MS spectrum of *m/z* 583.2382 at 19.9 min in the chromatogram of the sample taken after 24 h of incubation. MM-CAD, monomethylcadaverine; MML, monomethyllysine; Subscript ds indicates dansylated products. For DML: (d) RP-HPLC with UV detection and (e) MS/MS detection after dansylation. (f) MS/MS spectrum of *m/z* 364.2050 at 9 min in the chromatogram of the sample taken after 24 h of incubation. For ease of comparison, this panel replicates the spectrum shown in Figure 2c of the main manuscript. Abbreviations: DM-CAD, dimethylcadaverine; DML, dimethyllysine.

**Figure S12.**
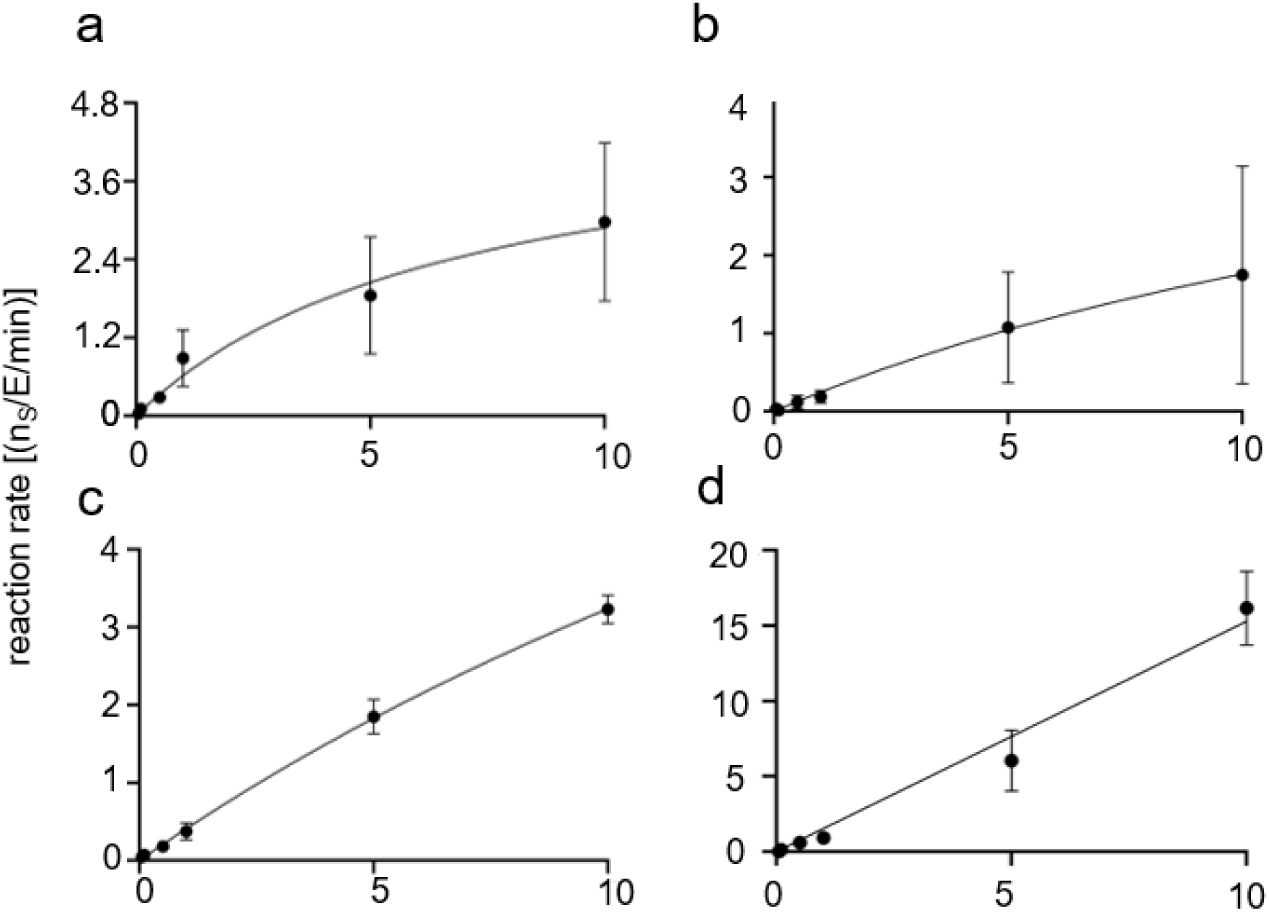
Kinetic curve for SpeC with (a) FmL, (b) MML, (c) DML and (d) lysine as substrate. The enzyme concentration was 500 nM and the substrate concentration ranges from 0.05 to 10 mM. The reaction product was quantified by HPLC-UV after pre-column dansylation. For ease of comparison within this extended dataset, the kinetic curve of FmL (panel a) replicates the data shown in Figure 2f of the main manuscript. The results are represented as mean and standard deviation of at least three independent replicates.

**Figure S13.**
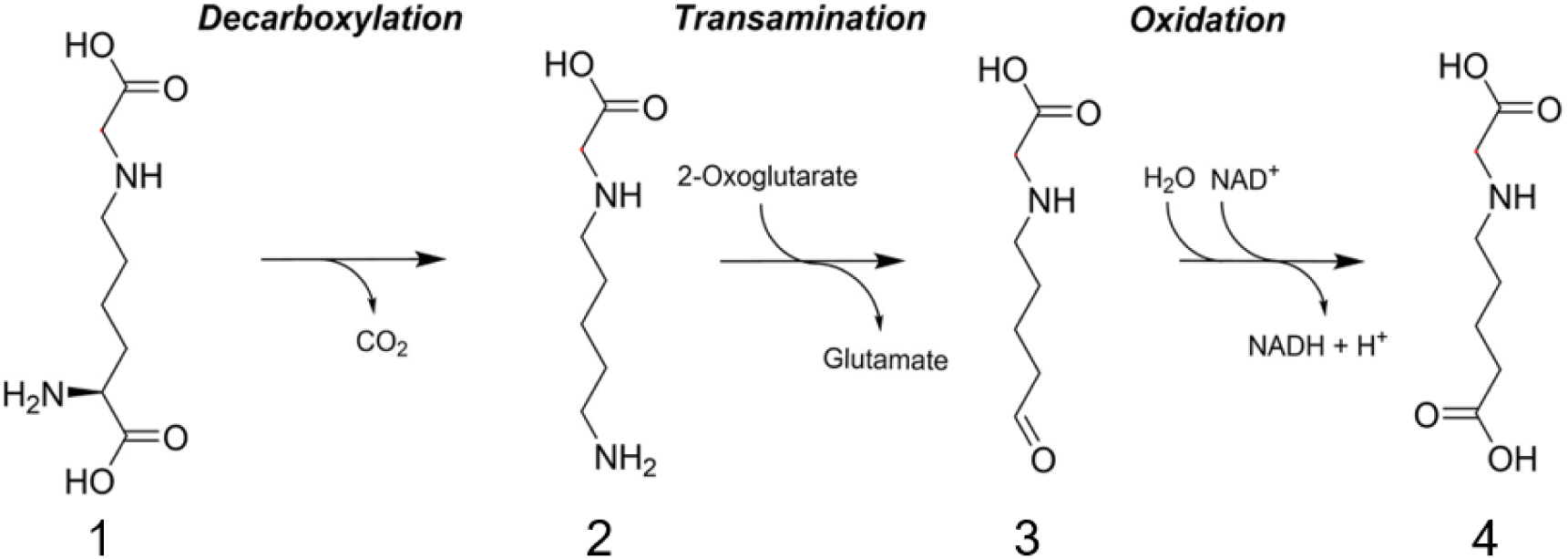
Proposed degradation pathway based on the metabolites formed during the incubation of CML with *E. coli.* 1) CML 2) CM-Cad, 3) carboxymethylaminopentanal, and 4) carboxymethylaminopentanoic acid (CM-APA). Figure modified from previous literature (Mehler et al., 2022).

**Figure S14.**
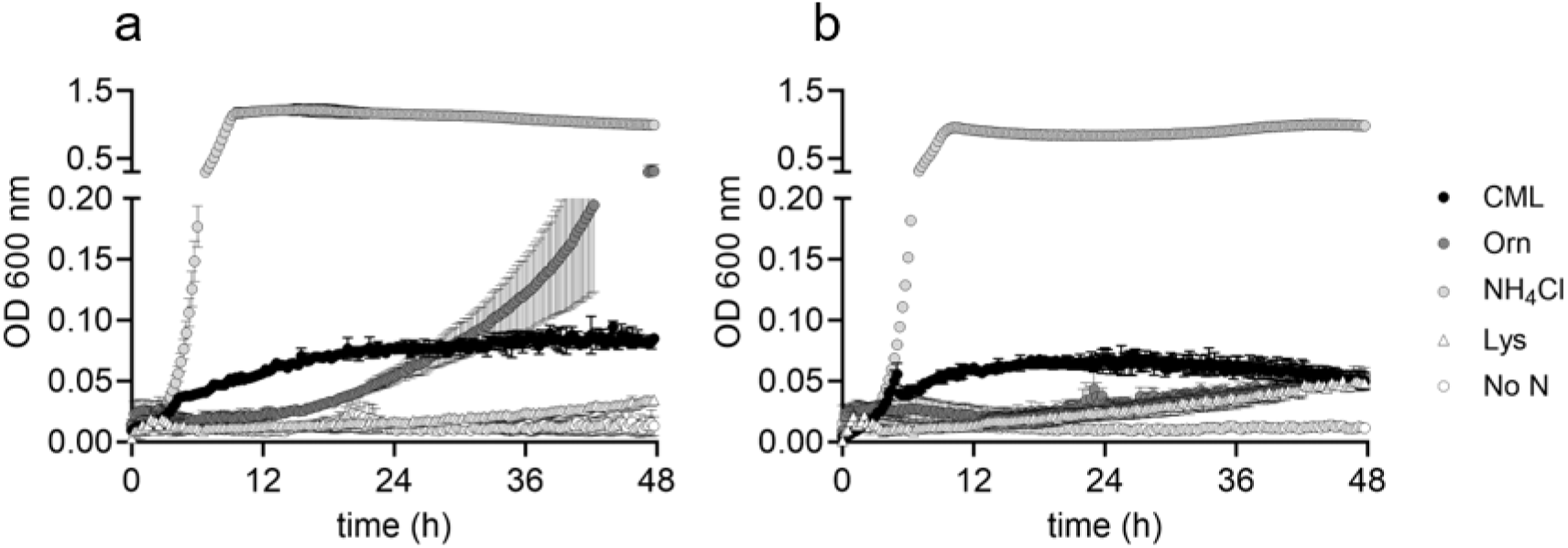
Growth of *E. coli* BW25113 (a) and the mutant Δ*speC* (b) in M9 minimal medium in presence of different nitrogen sources in 10 mM concentration. The OD 600 nm was recorded spectrophotometrically for 48 h. The values are shown as mean and standard deviation of three independent replicates.

**Figure S15.**
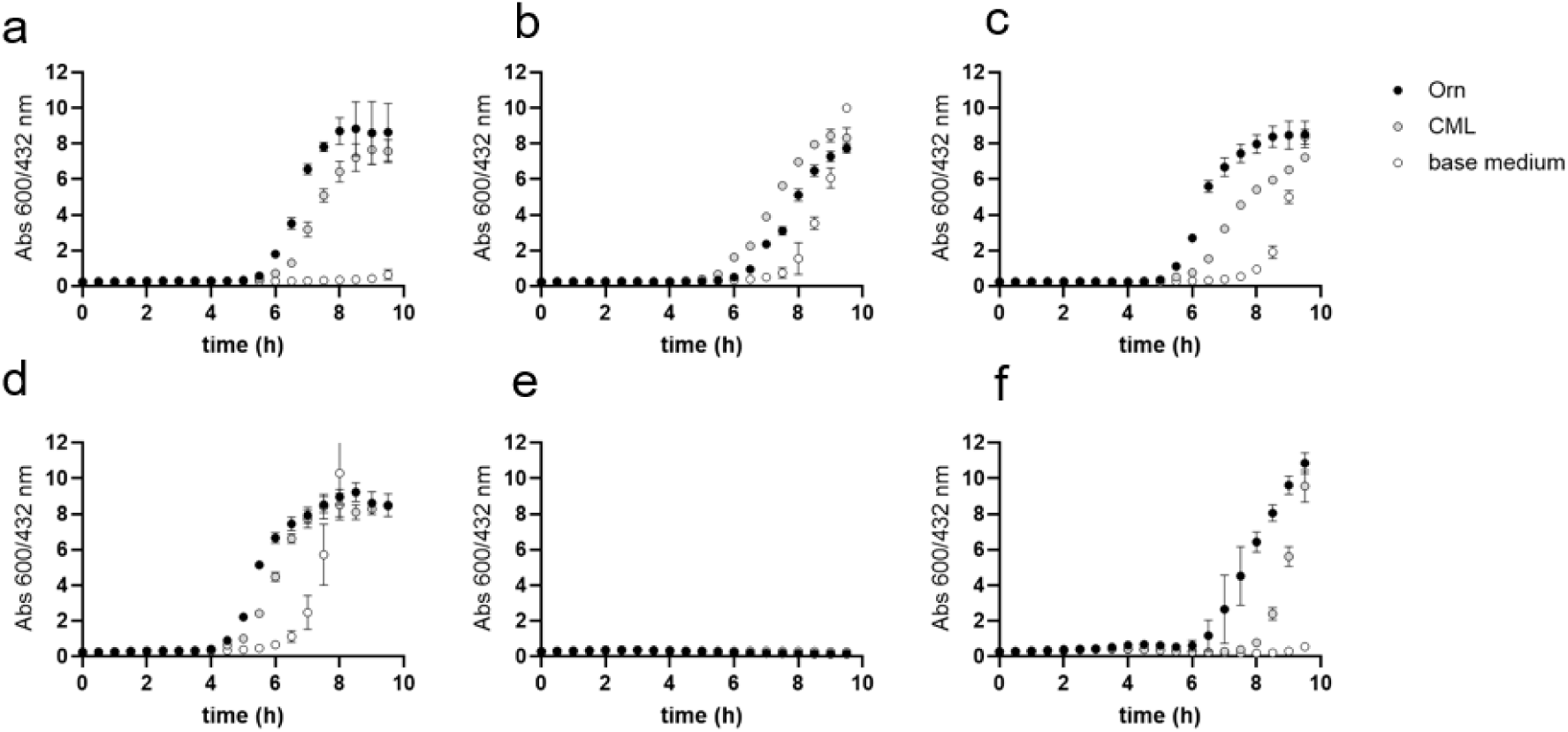
CML-dependent activation of the *E. coli* pH-stress response. Investigated was the ability to increase external pH by decarboxylation of ornithine (Orn) or CML in six strains of clinical relevance: O157 H7 (a), ATCC 8739 (b), O111 H- (c), UTI 89 (d), O127 H6 (e), CFT 073 (f). The restoration of neutral pH was monitored over 10 h through absorbance at 600 and 432 nm of the indicator bromocresol purple that was present in the medium adjusted to pH 5.0. A rise in the absorbance ratio to saturation indicates pH restored to neutrality. The values are shown as mean and standard deviation of three independent replicates.

**Figure S16.**
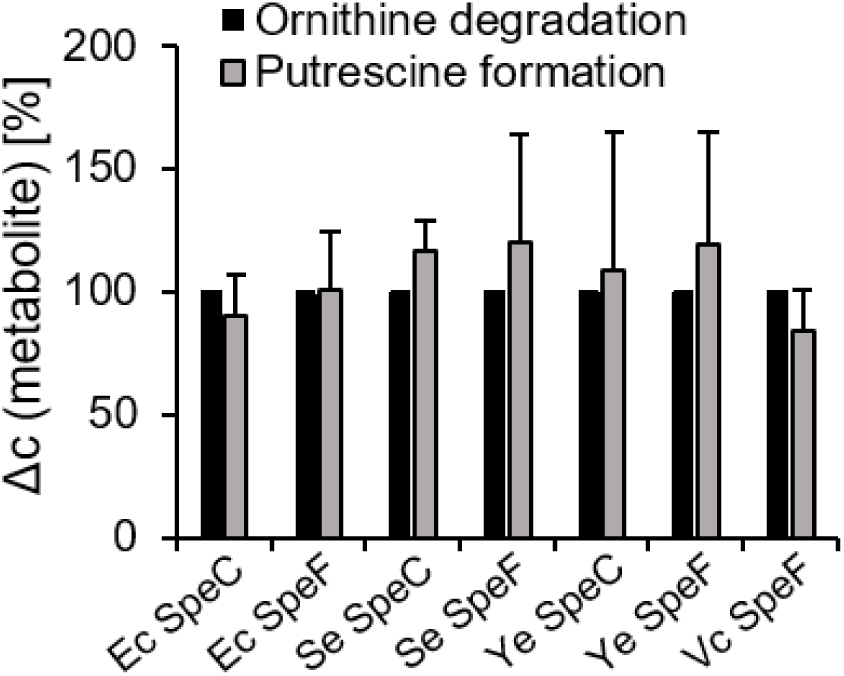
*In vitro* degradation of ornithine mediated by the decarboxylases SpeC and SpeF purified from: *E. coli*, *S. enterica*, *Y. enterocolitica* and *V. cholerae.* The metabolites were measured by HPLC-MS/MS and the amount of ornithine degraded and resulting putrescine formed are shown as percentage. The substrate was applied in a concentration of 10 mM, and the enzymes were applied in a concentration of 50 nM. Abbreviations: *Escherichia coli*, Ec; *Salmonella enterica*, Se; *Yersinia enterocolitica*, Ye; *Vibrio cholerae*, Vc. The values are shown as mean and standard deviation of three independent replicates.

**Figure S17.**
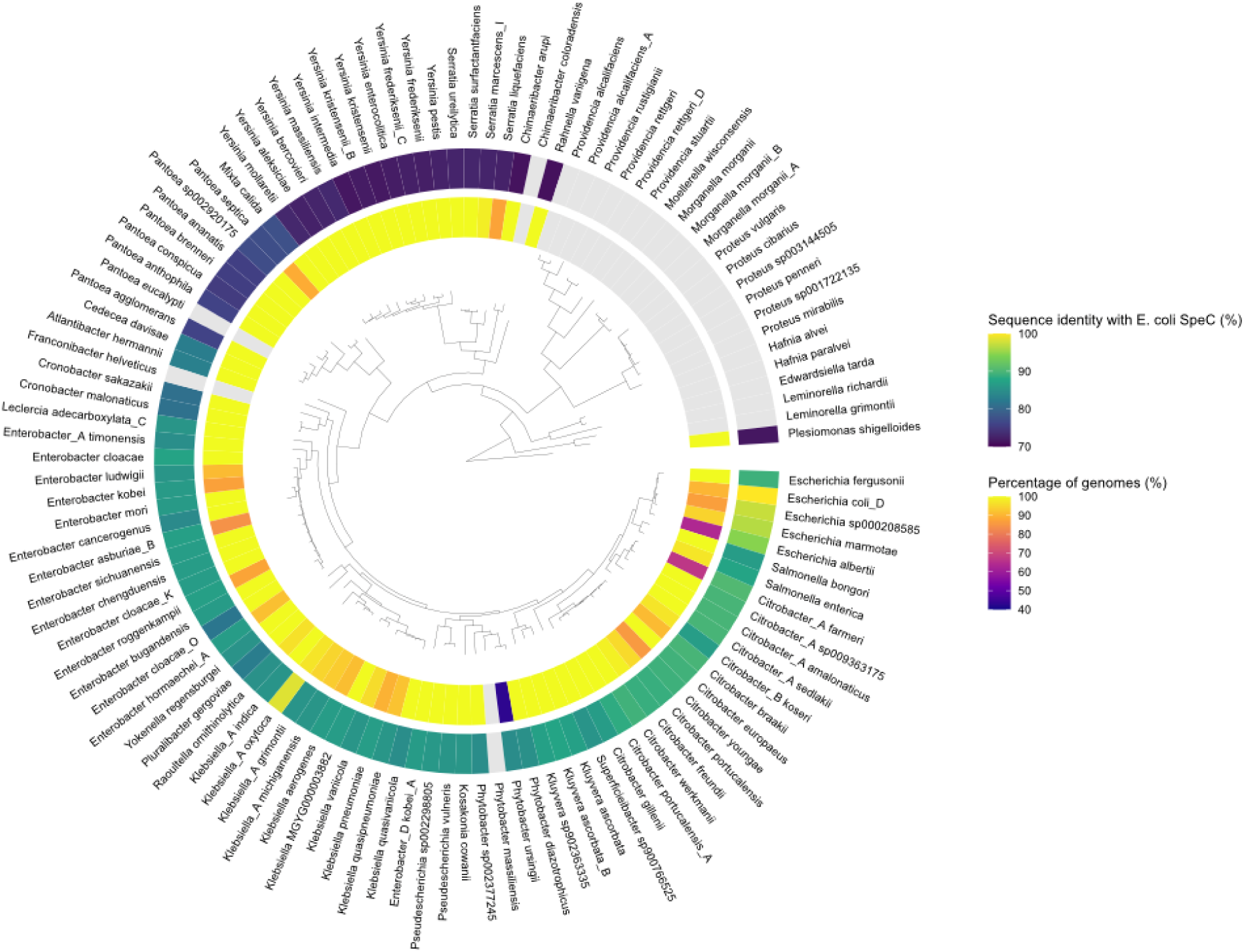
Occurrence of CML-degrading bacteria in the human gut microbiota. Using the Unified Human Gastrointestinal Genome (UHGG) and Protein (UHGP) collections (Almeida et al., 2021), we identified 10,252 genomes from 89 species belonging to 29 genera, in the human gut microbiota which are capable of CML degradation. The species in the Enterobacteriaceae family are shown. The outer ring shows a heatmap of the sequence identity between *speC* orthologs in each species and E. coli *speC*. A CML degradation capability is defined as the presence of an ortholog of *speC* which has a sequence identity that is at least 70% with the *E. coli speC*. The inner ring shows a heatmap of the percentage of genomes capable of degrading CML in each species. Grey indicates no genome in a species was identified as CML-degrading.

**Figure S18.**
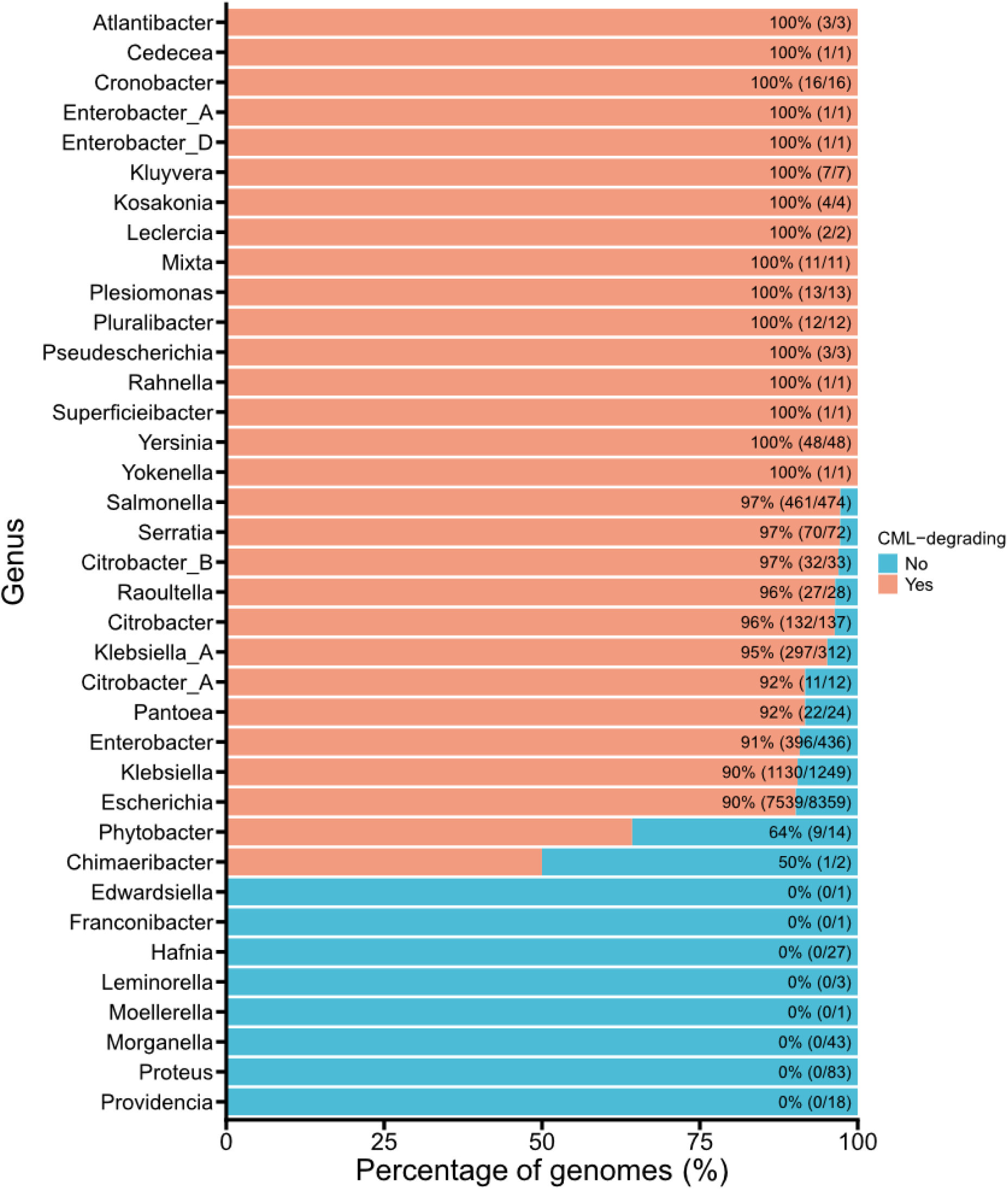
Occurrence of CML degrading bacteria in the 29 Enterobacteriaceae genera, based on the presence of a *speC* ortholog whose sequence identity is at least 70% with the gene of *E. coli*.

**Figure S19.**
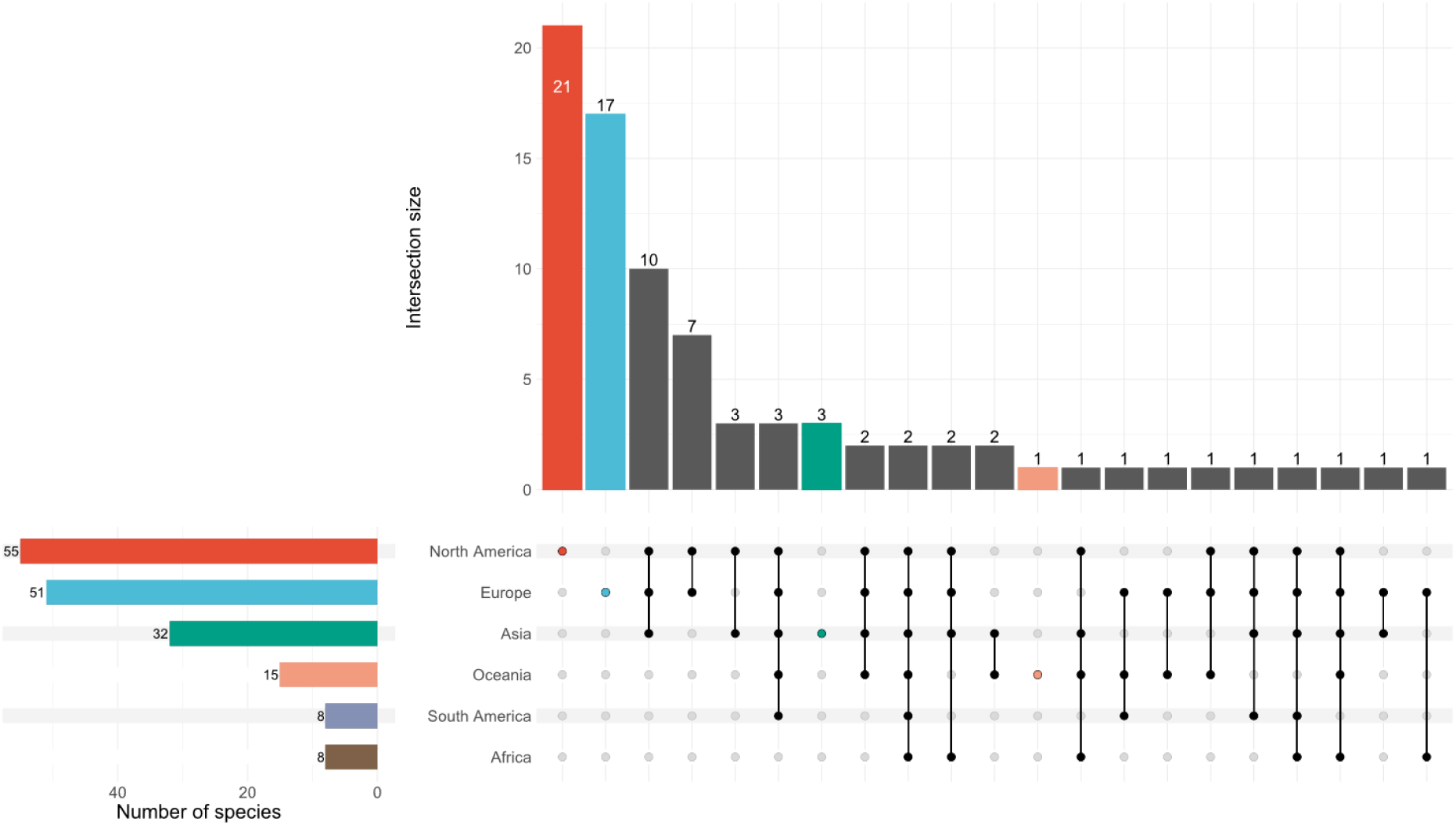
Number of species identified as CML-degrading across continents, ordered by their level of overlap. Vertical bars indicate the number of species shared between the continents highlighted with colored dots in the lower panel. Horizontal bars in the lower panel indicate the total number of species contained in each continent.

**Figure S20.**
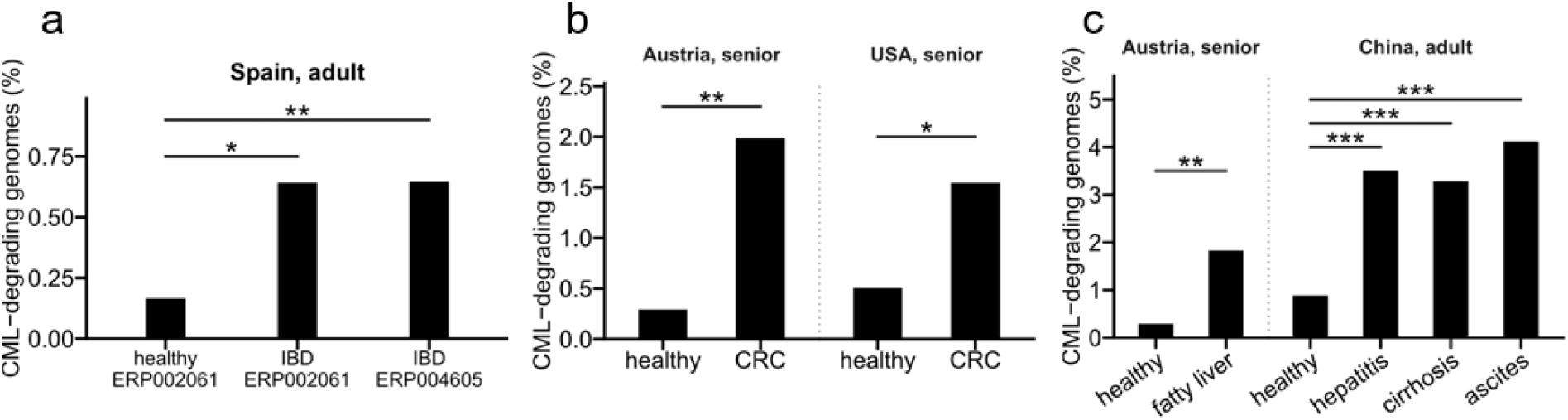
(a) Occurrence of CML-degrading bacteria in the gut microbiota in adults with IBD from Spain (Studies ERP002061 and ERP004605). *P-*values of the Fisher’s Exact test are: 0.00873 (top) and 0.0154 (bottom). (b) Occurrence of CML-degrading bacteria in the gut microbiota in senior people affected by CRC from Austria and USA (Study ERP008729) and the USA (Study ERP013933). *P-*values of the Fisher’s Exact test are: 0.00102 (left) and 0.0121 (right). (c) Occurrence of CML-degrading bacteria in the gut microbiota of senior people affected by fatty liver from Austria (Study ERP008729) and hepatitis, cirrhosis or ascites from China (Study ERP005860). *P-*values of the Fisher’s Exact test are: 0.00158 (left), 6.51e-8 (top-right), 3.94e-7 (middle-right), 2.62e-7(bottom-right).

